# A new typing scheme demonstrates high discriminatory power for *Treponema pallidum* subspecies

**DOI:** 10.1101/2025.07.10.664125

**Authors:** Marta Pla-Diaz, Lorenzo Giacani, Lauren C. Tantalo, Mahashweta Bose, Tara B. Reid, Christina M. Marra, David Šmajs, Petra Pospíšilová, Klára Janečková, Takuya Kawahata, Fumiya Banno, Kendra Vilfort, Weiping Cao, Allan Pillay, Angel Noda, Philipp P. Bosshard, Marcus Chen, Oriol Mitjà, Verena J. Schuenemann, Simon Hackl, Kay Nieselt, Pablo Hernández-Bel, Ma Dolores Ocete, Natasha Arora, Fernando González-Candelas

## Abstract

The global resurgence of treponematoses, particularly syphilis, poses a growing public health challenge. Despite recent advances in sequencing technologies, obtaining complete *Treponema pallidum* genome sequences for epidemiological studies remains time-consuming and challenging due to the difficulty related to procuring clinical samples with sufficient treponemal burden to fulfil the sequencing requirements. There is an urgent need for rapid, cost-effective and accessible typing methods suitable for laboratories with Sanger sequencing resources. Based on the analysis of 121 *T. pallidum* genomes from geographically diverse regions, we selected seven highly variable genes to form the basis of this new typing system. These seven genes show high discrimination capacity, identifying many allelic profiles among *T. pallidum* isolates. Importantly, the scheme employs a single-step PCR protocol for the amplification and sequencing of all seven targets enabling straightforward implementation in standard laboratory settings. The MLST was validated using a diverse set of *T. pallidum* clinical samples from across the globe. A significant proportion of the tested samples showed macrolide resistance, emphasizing the need for epidemiological surveillance. Utilizing this new tool, we have analyzed the genetic variation within and between populations of *T. pallidum*, considering the geographical origin of the samples. Population structure analysis revealed distinct genetic clusters, underlining complex transmission dynamics of *T. pallidum*, shaped by local epidemiological factors. The MLST scheme is publicly accessible through the PubMLST database, encouraging widespread adoption in standard laboratories due to this database being user-friendly, intuitive, and fast to implement. The novel MLST scheme offers a promising tool to advance the study of the molecular epidemiology of *T. pallidum*, facilitate tracking transmission, and establish a global surveillance network with the overall goal of strengthening public health interventions for syphilis control.

## 1. INTRODUCTION

*Treponema pallidum* subsp. *pallidum* (TPA) is an extracellular gram-negative bacterium and the causative agent of syphilis, a multistage and mostly sexually transmitted infection (STI). There are similar diseases called endemic treponematoses, namely yaws, bejel and pinta, caused by closely related bacteria: *T. pallidum* subsp. *pertenue* (TPE), *T. pallidum* subsp. *endemicum* (TEN), and *T. carateum*, respectively. Until very recently, the transmission of endemic treponematoses was considered to occur through non-sexual skin contact. However, recent studies from France, Cuba and Japan have identified TEN in genital lesions among sexually-active males, supporting transmission through sexual contact [1–3].

The incidence of syphilis continues to increase worldwide, with more than 7 million cases reported annually in 2020 [4]. In 1997, the World Health Organization (WHO) estimated that more than 2.5 million people were infected with the endemic treponematoses [5]. Yaws was targeted by the WHO for eradication by 2020 through large-scale mass-treatment programs of endemic communities, but the disease is still prevalent in some regions [6–9].

All human treponematoses are multi-stage infections with variable clinical manifestations that can make accurate diagnosis difficult. Moreover, serological methods are unable to differentiate these diseases from each other [10]. With the introduction of molecular tools, it is now possible to identify all *T. pallidum* subspecies using genotyping and DNA-sequencing-based methods [1,11–19]. Despite these advances, cultivating these bacteria in the laboratory directly from clinical samples remains very challenging [20][21] and obtaining genomic sequences from clinical specimens requires expensive and time-consuming enrichment and sequencing techniques. These challenges highlight the need for more accessible genotyping tools to expand our understanding of the epidemiology of these bacteria.

Multilocus Sequence Typing (MLST) is a molecular technique that allows the characterization of bacterial isolates using the variable sequences of usually six or seven housekeeping genes [22]. Most bacterial species have sufficient variation in these genes to yield multiple alleles, allowing many distinct allelic profiles to be distinguished in many bacterial taxa. As a result, there are many standard MLST schemes available for different pathogens of epidemiological interest such as *Neisseria* spp. [22], *Staphylococcus aureus* [23], *Campylobacter jejuni* [24], or *Streptococcus pneumoniae* [25]. The *S. aureus* MLST method amplifies short fragments (450-500 bp) in seven highly variable housekeeping genes that have been successfully used to differentiate many strains [23].

Several typing schemes have been proposed for *T. pallidum* (Table 1) and have been applied to our current understanding of the molecular epidemiology of these treponematoses (citations showing application of these other tools). While much has been gleaned by employing these typing methods, they often pose technical challenges related to amplifying targets from clinical specimens with minimal *T. pallidum* DNA. In addition, concerns about some of these typing schemes have been raised due to potential intra-strain variability at the loci (*arp* and *tpr* genes) employed [11,26,27]. Furthermore, the existing MLST schemes (Table 1) were designed using limited genome information to type and characterize only the *pallidum* subspecies of this pathogen. The continued use of MLST schemes for molecular epidemiology purposes is required to limit the reliance of technical skills while ensuring high sensitivity and reproducibility, minimal costs, and easy accessibility of data through public databases[28]. Currently, only TPA allelic profiles are available publicly in PubMLST [12].

**Table 1.** Molecular typing schemes for *Treponema pallidum*. For each of the typing schemes listed, the target subspecies, loci typed and publication are provided.

| Molecular typing scheme | Target | Loci | Reference |
| --- | --- | --- | --- |
| CDC | TPA | *arp*, *tprE*, *tprG*, and *tprJ* + *rpsA* | [11,26] |
| SMBT | TPA | *tp0136*, *tp0548* and 23S rDNA | [29,30] |
| ECDCT | TPA | Original CDC typing scheme + *tp0548* | [15] |
| MLST | TPA | *tp0136*, *tp0548*, and *tp0705* + 23S rDNA | [12] |
| MLST | TPE | *tp0548*, *tp0136* and *tp0326* | [16] |
| MLST | TPE | *tp0488*, and *tp0548* | [31] |
| CDC_TPE | TPE | CDC and ECDCT typing schemes + *tp0279* | [17] |
| MLST | TEN | *tp0136*, *tp0326*, *tp0367*, *tp0488*, *tp0548*, *tp0859*, *tp0861*, *tp0865* and 16S rDNA + 23S rDNA | [1] |
| MLST | TPE | *tp0488*, *tp0548* and *tp0858* | [32] |

Herein, we used 121 *T. pallidum* genomes collected from across the globe to design a new MLST scheme that allows differentiation between the three subspecies of *T. pallidum* and, also, within lineages of TPA. Our proposed scheme will aid in identifying genetic diversity and transmission patterns for the three diseases, as well as establishing a global network for surveillance and investigation of possible outbreaks.

## 2. MATERIAL AND METHODS

A summary of the in silico workflow used to design the MLST scheme utilizing 121 *T. pallidum* genomes is shown in Figure 1.

**Figure 1.**
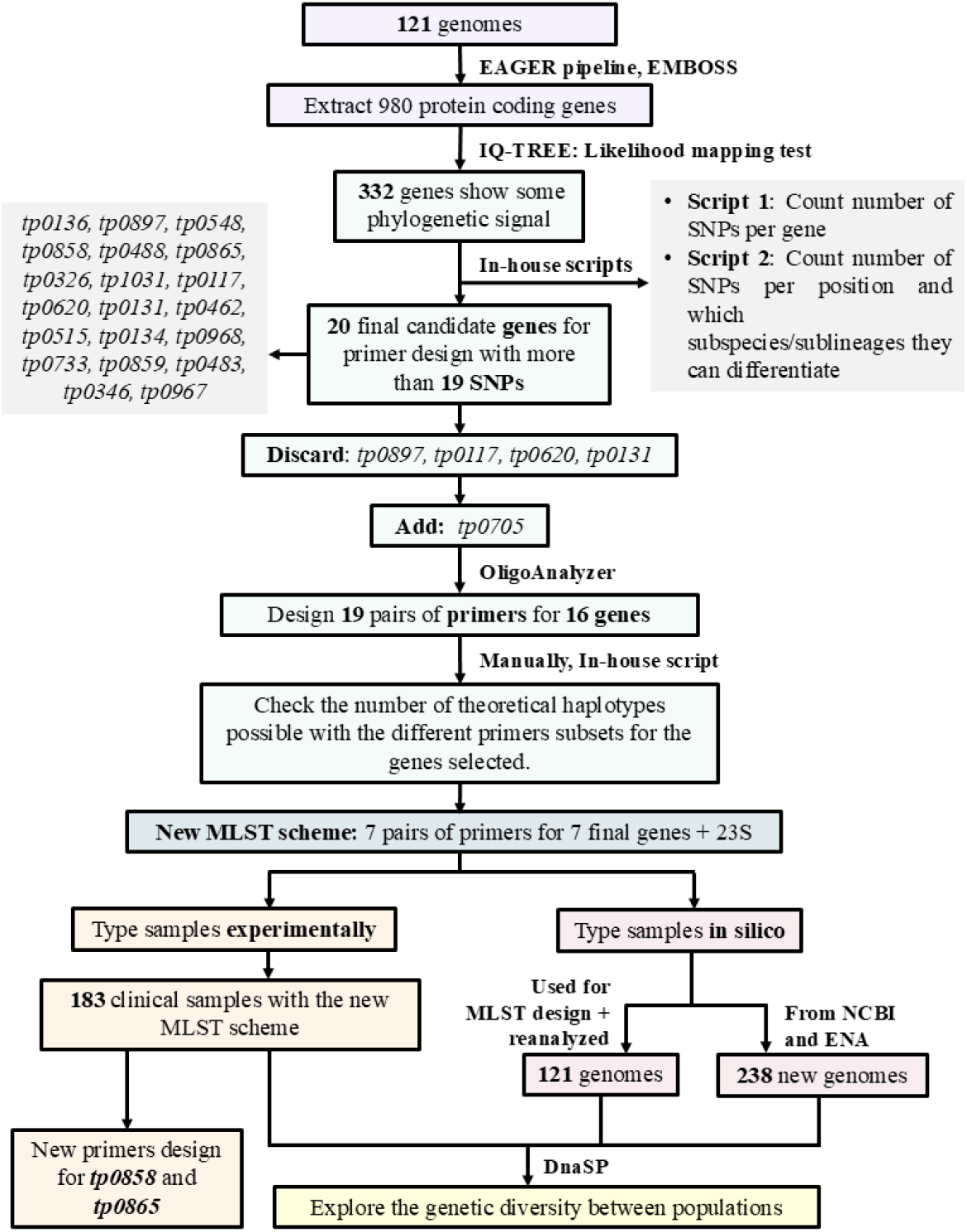
Workflow for the *in silico* and experimental design of a new MLST scheme.

### Genomic dataset generation and MLST design

We compiled a dataset with 121 *T. pallidum* genomes (Supplementary Table 1A). The dataset included 101 TPA genomes (14 from the Nichols clade and 87 from the SS14 clade), 17 TPE genomes and 3 TEN genomes from previous studies and public databases accessed up until December 2018. This dataset included 26 newly sequenced genomes (now available at Supp. Table 1A and BioProject PRJNA1288478).

For genomes with short read data available (n = 103), we used the EAGER pipeline [33] to reconstruct the individual genomes. For the few genomes with only consensus sequences available (n = 18), high throughput sequencing (HTS)-like reads based on genome assemblies were simulated using Genome2Reads [33]. After adapter clipping, merging and quality trimming, the resulting reads for each genome were mapped to the Nichols genome (NC_021490.2), using BWA-MEM [34] applying default parameters. PCR duplicates were removed with DeDUP [33]. QualiMap 2.17 was used to calculate the coverage breadth of the reference genome and coverage depth [35]. SNP calling was performed using GATK UnifiedHaplotyper 3.6 [36]. Genome sequences were excluded for further analysis if they did not have at least 80% of the Nichols genome covered with at least 3 reads per base [37]. Variant alleles were called if supported by at least 3 reads and a minimum frequency of 0.9. MUSIAL (https://github.com/Integrative-Transcriptomics/MUSIAL) was applied to obtain a multiple genome alignment (MSA) from the resulting VCF files.

### PCR target selection and design of pilot MLST method

We identified candidate loci for the new typing system using the information from 121 *T. pallidum* genomes. First, we assessed the phylogenetic information of each protein-coding gene using the likelihood mapping (LM) test as implemented in IQ-TREE [38], following the annotation of the Nichols reference genome. Genes with some phylogenetic signal, evaluated as likelihoods falling outside the central region in the LM triangle [39], were retained for the ensuing downstream analyses. Next, we checked the total number of single nucleotide polymorphisms (SNPs) per gene and which subspecies and/or sublineages could be differentiated by each of these SNPs (the scripts used are provided in Supplementary Files 1 and 2). Those genes with the highest level of variation and discriminatory power were selected as candidate MLST markers. For each of these genes, primers were designed following the criteria listed below:

1. Primers should anneal to conserved regions.
2. The 3’-end nucleotide should correspond to a second codon position of a coding sequence.
3. Primer length should be between 18 - 22 bp.
4. Amplicon size should be between 400 - 700 bp.
5. Primers should have 45%-60% GC content.
6. Forward and reverse primers should not be complementary.
7. Primers should have a minimum dG value of -6 kcal/mol.

Next, we prioritized primer combinations to maximize the number of haplotypes distinguished across the 121 genomes, with primer combinations designed for the selected genes (the script used can be found in Supplementary File 3). Finally, we selected the set of primers for seven genes yielding the highest level of resolution and tested them experimentally using a serial dilution (10^−1^ - 10^−8^) of the TPA Nichols’s strain DNA plus a negative control. The initial concentration of the Nichols DNA sample was determined to be 34.5 ng/μL, corresponding to 103 ng per PCR reaction.

The primers used for amplification of the macrolide resistance markers in the 23S rRNA amplification were the same as those previously published by Lukehart *et al*. [40].

### Clinical sample typing: sample collection, DNA extraction, and sequencing

We obtained 179 clinical samples of *T. pallidum* (Table 2) to be typed with the new MLST scheme using Sanger sequencing.

**Table 2.** Clinical samples collected to be typed by Sanger sequencing with the new MLST scheme. The table shows the number of samples obtained for each of the subspecies of *T. pallidum*, as well as the group providing the samples (source) and the location at which the DNA extraction and typing of the samples with the new MLST scheme was conducted. (CHGUV: Consorcio Hospital General de València; FISABIO-UV: FISABIO-University of Valencia; MU: Masaryk University; UW: University of Washington; CDC: Centers for Disease Control and Prevention; OIPH: Osaka Institute of Public Health).

| Subspecies | Number of samples | Source | DNA extraction | MLST tests |
| --- | --- | --- | --- | --- |
| TPA | 76 | CHGUV | FISABIO-UV | FISABIO-UV |
|  | 21 | MU | MU | MU |
|  | 36 | UW | UW | UW |
| TPE | 16 | MU | MU | FISABIO-UV |
|  | 13 | UW | UW | UW |
|  | 8 | CDC | CDC | CDC |
| TEN | 5 | OIPH | OIPH | OIPH |
|  | 4 | MU | MU | MU |

In addition, four historical isolates were obtained from the McGovern Medical School and propagated in rabbits as previously described for subsequent DNA isolation and typing at FISABIO-UV. These included three TPE isolates (Gauthier, Samoa D, and Samoa F) and the Nichols TPA isolate.

The seven loci (*tp0136, tp0326, tp0548, tp0705, tp0858, tp0865* and *tp1031*) selected using the methods described earlier and the 23S rRNA gene were amplified by PCR with a touchdown protocol. The total volume of each of the seven PCR reactions was 25 μL, with the following composition: 3 μL of DNA, 1.5 μL of dNTP mix from TaKaRa Ex Taq® DNA Polymerase 250 Units kit, 2.5 μL Mg2+ plus buffer and 0.05 μL Taq polymerase, 0.25 µL of each primer (10 µM), and 0.75 μL of DMSO (3%) to increase the specificity and yield of the PCR reaction.

Touchdown PCRs consisted of 40 amplification cycles. In the first 10 cycles, we used a melting temperature 4ºC higher than the optimal melting temperature to increase specificity. For the remaining 30 cycles, we used the predicted optimum melting temperature for each primer.

The seven PCR products were purified using the NucleoFast 96 PCR Plate (Cultek). PCR reactions were adjusted to 100 µL with milliQ water and transferred to the purification plate. Samples were centrifuged at 3500 rpm for 10 min, washed with 100 µL of milliQ water and centrifuged again under the same conditions. A final elution was performed by adding 30 µL of milliQ water to each well, followed by incubation for 10 minutes at room temperature with gentle agitation (300 rpm). Eluted DNA was collected, transferred to a new plate, and stored at −20 °C. Purified products were afterwards sequenced by Sanger sequencing.

Sequence analyses were performed using the Staden package v4.11.2-r [41] and merged in a gene alignment for each locus to verify the quality of each sequence and obtain its corresponding allele and sequence type (ST) by inspection with AliView 1.25 [42].

For the 23S rRNA gene, positions 2058 and 2059 were assessed for A→G mutations indicative of macrolide resistance [43,44].

### Validation of the new MLST scheme

We applied the newly developed MLST scheme to three types of data. First, we analyzed sequencing read data obtained directly from clinical and historical samples (n = 183), as previously described. Second, we reanalyzed the 121 genomes originally used in the development of the scheme, which were also available as read data. Finally, we applied the scheme *in silico* to all *T. pallidum* genome assemblies publicly available in the European Nucleotide Archive (ENA) and the National Center for Biotechnology Information (NCBI) as of September 2022 (n = 238). Of these, 226 were TPA, 5 TPE, and 7 TEN genome assemblies (see Supplementary Table 1B for details).

All 238 genome assemblies were consolidated into a single FASTA file and aligned using MAFFT v7.467 [45] to generate a whole-genome alignment. From this alignment, the regions flanking the primer binding sites of each locus in the MLST scheme were extracted, producing eight separate FASTA files, one per locus. To assign allelic profiles, CD-HIT v4.7 [46] was used to cluster sequences based on similarity. Cluster assignments were manually verified using AliView v1.25 [42] to ensure accuracy. Sequence comparisons were performed to identify allelic differences, including single nucleotide polymorphisms (SNPs) and insertions/deletions (indels), which were then used to define distinct alleles. The combination of alleles across the seven loci determined the sequence type (ST) for each genome.

In light of recent evidence suggesting that reference genome selection can introduce biases in variant calling and downstream analyses [47], we mapped and processed the 121 genomes used in the original MLST scheme design against three additional reference genomes selected as representative of distinct phylogenetic groups: SS14 (NC_010741.1) for the SS14-like clade within TPA, CDC2 (NC_016848.1) for TPE, and BosniaA (NZ_CP007548.1) for TEN, in addition to the original Nichols reference for Nichols-like TPA strains. For each genome, we extracted the gene sequences corresponding to the seven MLST loci from the realignment that best matched the corresponding subclade or subspecies. These final sequences, minimizing potential reference bias, were used for allele assignment and sequence type (ST) designation according to the MLST scheme.

### Genetic diversity and population divergence in *T. pallidum*

We compared the genetic diversity (D) of STs between the different subspecies/sublineages of *T. pallidum* using the expression D = l-∑p_i_ ^2^ [48] where *p*_*i*_ is the frequency of each ST for each subspecies/sublineages of *T. pallidum*.

Additionally, to analyze the geographical distribution of the different STs, we examined the genetic diversity within and differentiation between populations using DnaSP 6.12.03 [49]. The populations referred to either the countries or the continents of origin of the samples. We excluded Oceania as it only had two samples. Due to the small sample size available for the TPE and TEN subspecies, we performed these analyses only for TPA. Next, at both the continental and country level, we estimated the average number of nucleotide differences (*k*) within and between populations. Moreover, we also estimated the nucleotide diversity (П) to examine the degree of polymorphism within populations, and between populations using the Jukes-Cantor model (Da(JC)) of nucleotide substitution.

### Inference of phylogenetic trees

We reconstructed a maximum likelihood phylogenetic tree using the concatenated sequences of the seven loci for each of the samples for which an ST could be derived. For this, we used IQ-TREE2 [38] with GTR+G as the evolutionary model and 1000 bootstrap resampling replicates.

## 3. RESULTS

### Reference-based alignment for the design of a new MLST scheme

The resulting multiple sequence alignment from the 121 genomes (Supplementary Table 1A) spanned a total of 1,139,633 bp and allowed us to identify 3,465 SNPs.

### Selection of loci

From the 978 protein-coding genes extracted using the Nichols genome as reference, a likelihood mapping test was performed for 780 genes, after excluding 198 genes due to the large number of undetermined positions. The analysis resulted in a total of 332 genes displaying some phylogenetic signal (Supplementary Table 2).

To prioritize subsets of genes to provide optimal resolution for the new MLST scheme, we assessed the number of SNPs in each of these 332 genes (Supplementary Table 3) and which subspecies and/or sublineages could be differentiated by each SNP. This procedure led to a selection of 20 candidate genes (Supplementary Figure 1) for primer design. These 20 candidate genes each contained 19-120 SNPs, displaying the largest variation and power of discrimination for the three different subspecies (Supplementary Table 4).

After excluding 5 genes (*tp0897, tp0117, tp0620, tp0131, tp0733)* that did not contain suitable flanking primer binding sites, we designed 19 different PCR primer pairs for 15 of the 20 candidate genes. We included an additional gene in subsequent analyses, *tp0705*, because this gene added to the discriminatory power of the MLST scheme designed for TPA [12].

After designing the primers, we analyzed the potential number of haplotypes that could be differentiated using different combinations of the primers developed for the 16 selected genes (Supplementary Table 5). Based on this analysis, we selected a set of seven primers that yielded the highest discriminatory power for the four main subspecies/sublineages of *T. pallidum* (TEN, TPE, TPA Nichols and TPA SS14). These seven sets of primers target *tp0136, tp0326, tp0548, tp0705, tp0858, tp0865*, and *tp1031* (Supplementary Table 6). The specific combination of alleles in these 7 loci defines the sequence type (ST) of each strain. This primer set also consistently amplified a serial dilution (10^−1^ - 10^−8^) of the Nichols DNA sample (Supplementary Figure 2). Our analyses yielded an estimated 12 and 16 STs for the Nichols and SS14 lineages of TPA, respectively, and 11 and 3 STs for the TPE and TEN subspecies. Detailed information on the estimated number of STs for the 121 genome dataset is shown in Supplementary Table 7. In addition, we also analyzed the 23S rRNA gene to detect the two A→G mutations indicative of macrolide resistance.

During the experimental testing of the new MLST scheme (see below), we noted non-specific amplification of loci *tp0865* and *tp0858* in some TEN samples. Although the regions used for primer design were conserved for these loci in the 121 genomes dataset, we designed alternative primers for these two targets.

For locus *tp0858*, a new set of primers were designed to flank the initial primer set (Supplementary Table 8). The new reverse primer is not inside the *tp0858* gene; it is in the intergenic flanking region of the 3’ end. The new forward primer is inside the *tp0858* gene and downstream of the 5’ region of the original forward primer. We tested the new primers for the *tp0858* gene in five TEN strains from Japan (Osaka-2017A, Osaka-2017B, Kyoto-2017, Osaka-2018, Osaka-2018B), and upon sequencing amplicons it was discovered that the lack of amplification with the original *tp0858* primers (Supplementary Figure 3) was due to a deletion of 63 bases in the 5’ end in those specimens.

For locus *tp0865*, it was not possible to design new primers flanking the previous ones while keeping the amplicon within the desired size. Hence, we designed new forward and reverse primers for the *tp0865* gene to test them in combination with the previous ones. The different combinations of the primers were selected according to the optimal melting temperature and amplicon size (Supplementary Table 9). All the combinations of the original and new primers successfully amplified Nichols DNA but none of the five TEN samples from Japan mentioned above.

### Allelic profiles identified with the new MLST scheme

We tested the new MLST scheme by performing Sanger sequencing on 179 clinical samples and 4 historical isolates (Supplementary Table 10). In addition, we performed *in silico* analyses on the genome sequences from another 238 genomes obtained from public databases. These analyses were complemented with the examination of 121 genomes that were utilized for the design of the new MLST scheme (Supplementary Table 1A). We included 12 sequences obtained from 10 isolates that had already been sequenced at least once (8 isolates) or twice (2 isolates). These sequences were used as controls to identify any sequencing artifact potentially resulting in different STs despite their common origin (Supplementary Note 1).

With the 542 genomes assessed with the new MLST scheme, we obtained multiple alleles per gene, as detailed in Table 3. The gene with the largest number of alleles was *tp0548* (n=53), whereas the gene with the fewest alleles was *tp1031* (n=6). The sequences of the representative alleles obtained in this study for each gene can be found in the Supplementary Files 4-10. The final allelic profiles obtained for each sample are detailed in Supplementary Table 10.

All the identified alleles and STs that did not contain missing data have been deposited in the PubMLST [50] database specific for this MLST. The remaining alleles will be deposited pending further verification.

**Table 3.** Allelic diversity across the new MLST genes. The table shows the number of alleles obtained for each gene of the new MLST scheme in clinical samples analyzed experimentally, by Sanger sequencing, and from WGS data analyzed *in silico*. The alleles with undetermined positions obtained from the *in silico* analysis are also detailed in the table, besides the number of alleles uploaded to PubMLST.

|  | N | *tp0136* | *tp0326* | *tp0548* | *tp0705* | *tp0858* | *tp0865* | *tp1031* |
| --- | --- | --- | --- | --- | --- | --- | --- | --- |
| **Sanger** | 183 | 8 | 17 | 30 | 6 | 18 | 9 | 5 |
| **WGS** | 359 | 8 | 16 | 24 | 1 | 14 | 12 | 1 |
| **All alleles obtained** | 542 | 16 | 33 | 53 | 7 | 32 | 21 | 6 |
| **Alleles with undetermined positions in WGS data** | - | 5 | 5 | 2 | 0 | 6 | 2 | 0 |
| **Alleles uploaded to PubMLST** | - | 11 | 28 | 51 | 7 | 26 | 19 | 6 |

### Sequence types (STs) identified among all typed samples

We were able to assign a ST for 415 samples out of the 542 samples analyzed (Supplementary Table 11): 82 from Sanger sequencing and 333 from the WGS dataset (Table 4), identifying a total of 82 different STs. We also looked for the presence of the two mutations in the 23S rRNA conferring macrolide resistance and found resistance alleles in 98 of 183 samples from the Sanger dataset (58 with complete ST), and in 101 of 313 samples from the WGS dataset (Table 4).

**Table 4.** ST and antibiotic resistance information across samples. Total number of samples for which a complete or partial ST was obtained. The information is specified for the Sanger sequence data from the clinical samples and for the WGS data analyzed *in silico*. The columns denote the number of samples with and without the mutation in the 23S rRNA conferring macrolide resistance, specified as resistant (R) or sensitive (S) as well as the number of samples for which the ST and 23S rRNA gene data could be combined.

|  | N | Complete ST | Partial ST | 23S (R) | 23S (S) | STs + 23S (R) | STs + 23S (S) |
| --- | --- | --- | --- | --- | --- | --- | --- |
| **Sanger** | 183 | 82 | 101 | 98 | 53 | 58 | 21 |
| **WGS** | 359 | 333 | 26 | 101 | 141 | 101 | 141 |
| **All** | 542 | 415 | 127 | 199 | 194 | 159 | 162 |

In addition, we analyzed the number of STs found within each subspecies of *T. pallidum*, and within each of the two lineages of TPA, as well as the number of samples resistant to macrolides (Table 5).

**Table 5.** STs obtained across *T. pallidum* subspecies and TPA lineages. The table shows, for each subspecies and sublineage, the number of samples available (N), the number of samples with and without the macrolide resistance mutation, specified as 23S S or 23S R, and the number of complete STs obtained. In addition, the table also shows the total number of different STs obtained and the genetic diversity for each subspecies/lineage.

| Subspecies/<br>sublineage | N | 23S (R) | 23S (S) | STs<br>Completed | STs + 23S | Partial<br>STs | Different<br>STs | ST<br>diversity<br>(D) |
| --- | --- | --- | --- | --- | --- | --- | --- | --- |
| **TPA-SS14** | 312 | 167 | 46 | 262 | 188 | 50 | 29 | 0.75 |
| **TPA-Nichols** | 126 | 12 | 102 | 120 | 110 | 6 | 31 | 0.88 |
| **TPA\*** | 19 | 11 | 1 | 0 | 0 | 19 | 0 | - |
| **all TPA** | 457 | 190 | 149 | 382 | 298 | 75 | 60 | 0.87 |
| **TPE** | 66 | 0 | 38 | 22 | 14 | 44 | 16 | 0.92 |
| **TEN** | 19 | 9 | 7 | 11 | 9 | 8 | 6 | 0.81 |
| **All** | 542 | 199 | 194 | 415 | 321 | 127 | 82 | 0.89 |

\* TPA samples not assigned to a specific TPA sublineage.

As illustrated in Table 5, the number of samples for which STs could be successfully determined differed across the subspecies and sublineages, reflecting the sample sizes in the dataset. For the TPA sublineages, complete STs were obtained for 382 samples (262/312 samples for SS14 and 120/126 for Nichols sublineage), while 22/66 samples and 11/19 samples yielded complete STs for TPE and TEN, respectively. As expected based on these sample sizes, the number of different STs present across the TPA sublineages was also higher than within TPE and TEN. We obtained 29 different STs for the TPA-SS14 lineage (Table 5): among these, ST22 was the most frequent, found in a total of 102 samples (39.1%). For TPA-Nichols, we obtained 31 different STs (Table 5) the most frequently found ST is ST57 (28.3%). We obtained 16 different STs for TPE (Table 5) and 6 for TEN (Table 5): the most frequent STs were ST12 (4/22 samples) for TPE, and ST42 (3/11 samples) for TEN.

We calculated the ST diversity (D) for the different subspecies of *T. pallidum*, using data on ST frequencies (Table 5). As observed in Table 5, TPE exhibited the greatest diversity (D=0.92), followed by TPA-Nichols (D=0.88), TEN (D=0.81), and finally TPA-SS14 (D=0.75).

Macrolide resistance alleles were detected in all the subspecies/sublineages (Table 5), except for TPE. TPA-SS14 was the sublineage with the largest number of resistant samples (167/312), compared to TPA-Nichols and TEN, with 12/126 and 9/19 resistant samples, respectively.

### Phylogenetic analysis

A maximum likelihood tree was constructed for the concatenated data of the seven loci included in the new MLST scheme (Figure 2).

**Figure 2.**
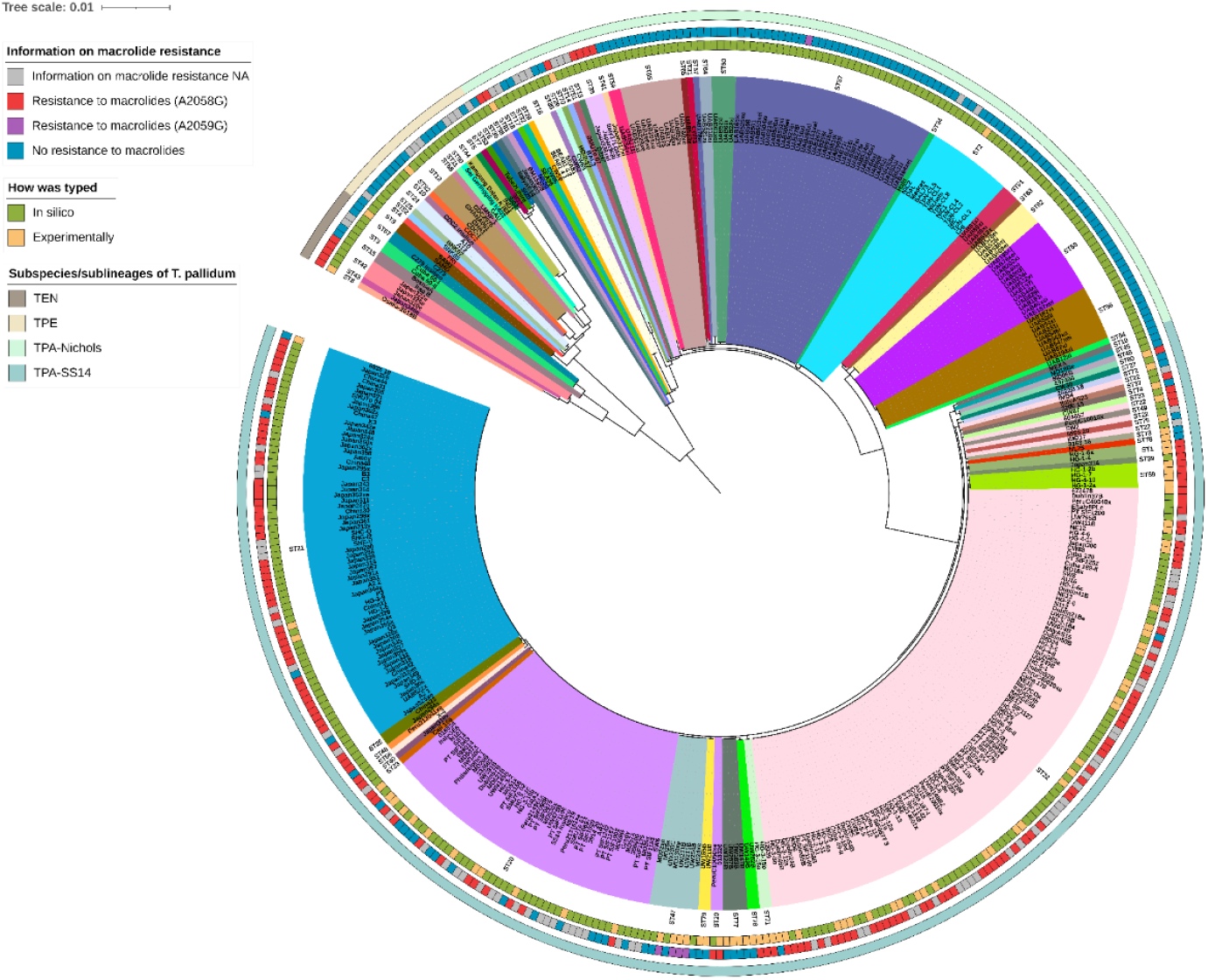
Maximum likelihood phylogenetic tree for the MLST data from 386 samples. The genetic data obtained through Sanger sequencing or WGS was concatenated and presented together with information on STs and macrolide resistance profile per sample.

The different STs assigned *in silico* or experimentally and the macrolide resistance mutations obtained in the 23S rRNA gene per sample are also shown in Figure 2. As observed, the four major clades corresponding to each of the *T. pallidum* subspecies and sublineages (TEN, TPE, SS14 and Nichols) were well supported in the maximum likelihood (ML) tree built from the concatenated alignment of the 7 MLST scheme loci. Notably, almost all the STs analyzed formed monophyletic clades, with the exceptions of three STs that formed paraphyletic groups (ST20, ST22 and ST57, Figure 2).

### Population genetic structure

We assessed the geographical distribution of STs identified with the new MLST scheme, focusing on the TPA subspecies, due to the limited availability of samples for TPE and TEN (Supplementary Table 11).

At the continental level (Supplementary Tables 12–13), the Americas showed the highest nucleotide diversity (k=15.958, π=0.00544). Asia (k=6.22, π=0.00210) and Africa (k=5.50, π=0.00197) showed intermediate within-group variation, while the lowest values were observed in Europe (k=5.29, π=0.00185). Notably, although Europe had the largest number of samples from multiple countries, it exhibited the lowest within-continent variation.

Overall, the average number of net nucleotide differences (k) between continents was larger than within continents. The highest intercontinental differentiation was observed between Africa and Asia (k=39.40, Da(JC)=0.00981), closely followed by Europe and Africa (k=39.16, Da(JC)=0.01023). The lowest differentiation was found between Europe and Asia (k=8.19, Da(JC)=0.00028). Oceania was excluded from these analyses due to the presence of only two samples.

At the country level (Supplementary Tables 14–15), Ireland and Portugal had the lowest within-country nucleotide diversity, with k=0.18 and 0.52, respectively. On the other hand, Italy and Cuba showed the highest values, with k=21.64 and 19.67, respectively, closely followed by the USA (k=17.88). The average number of net nucleotide differences between countries was higher than the average number of nucleotide differences within countries for Madagascar, Spain, Portugal, Ireland, Japan, China, Switzerland, and Czechia.

As illustrated in Figure 3, Madagascar is the country with the largest number of studied samples (n=85) followed by the USA (n=68). Both countries also have the highest numbers of different STs. A total of 14 different STs were found in the USA, the most frequent of which was ST20 (22/68 samples). In Madagascar, 13 different STs were observed, the most frequent being ST57 (34/85 samples).

**Figure 3.**
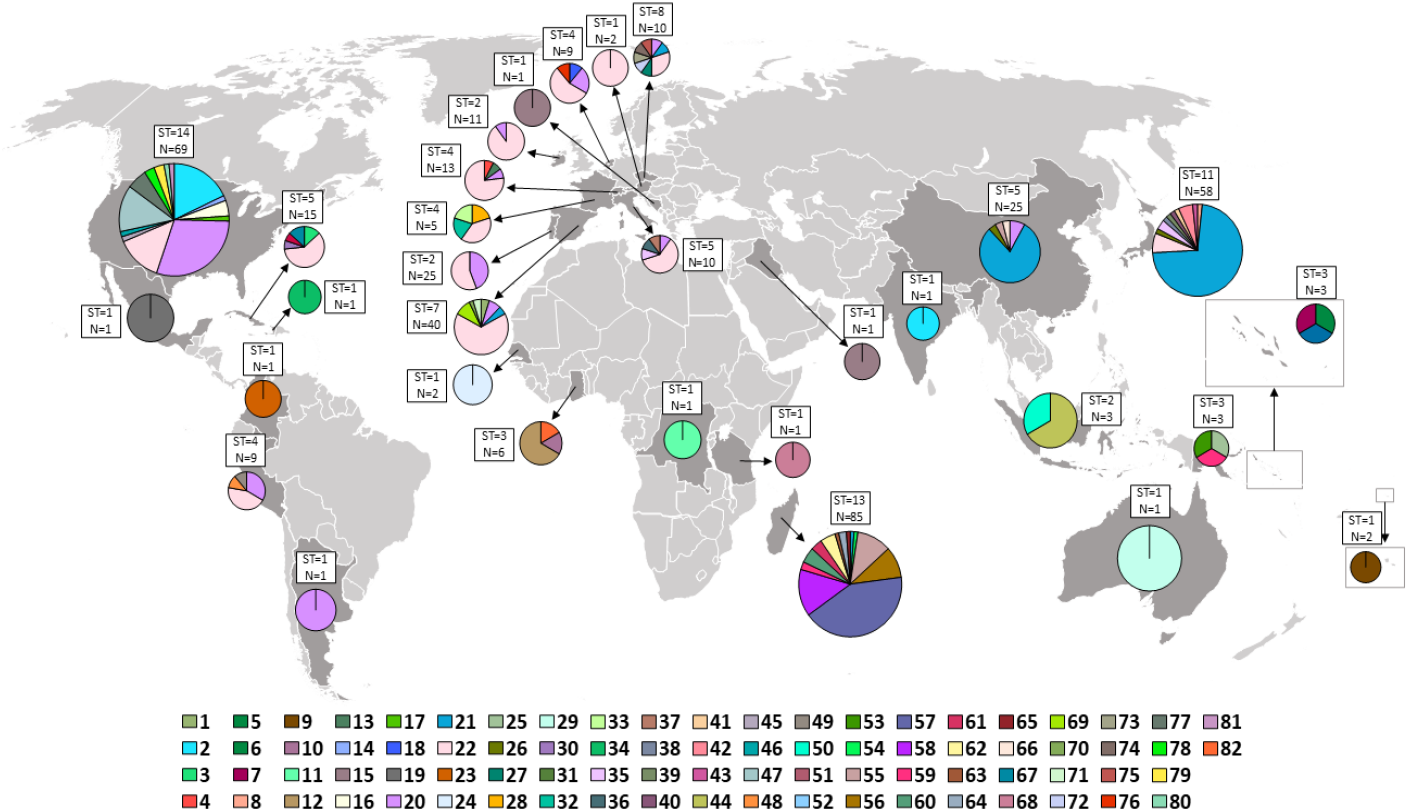
Geographical distribution of TPA STs identified with the new MLST scheme for *T. pallidum*. For each country, the number of samples and the number of different STs obtained is indicated. The legend shows the different colors for each identified ST.

## 4. DISCUSSION

In recent years, the incidence of human treponematoses, mainly syphilis, has increased markedly worldwide and has become a serious global health problem [1,2,4,6–9]. Currently, the only culture method for *T. pallidum* requires extensive infrastructure and expertise to isolate strains by rabbit propagation to reproducibly obtain high quality DNA for robust genome sequencing. Although the number of draft *T. pallidum* genomes available has increased since 2016 due to the introduction of enrichment techniques and WGS approaches to directly sequence *T. pallidum* from clinical samples [9,13,51–54], obtaining whole genome sequences is still a very time-consuming and costly process. For all these reasons, quick and economical typing procedures are desperately needed. Such a typing method can provide details on the frequency and incidence of strain types, longitudinal variation, associations with patient populations and communities, and antibiotic resistance trends. To maximize the information revealed by typing, we benefited from the recent increase in available *T. pallidum* whole genome sequences.

In this study, we analyzed 121 genome sequences of *T. pallidum* and developed an innovative and efficient molecular typing scheme. Our objective was to minimize the number of typing loci while remaining consistent with the standard number of markers in MLST schemes [22]. It is important to note that *T. pallidum* is considered a monomorphic bacterium and, therefore, each SNP can provide valuable information for strain characterization. Thus, we specifically searched for loci with the highest SNP densities to best capture overall diversity and to maximize the resolution, differentiation within each lineage and sublineage of this bacterium and distinguishing among the three *T. pallidum* subspecies investigated.

Our findings identified seven variable genes (*tp0136, tp0326, tp0548, tp0705, tp0858, tp0865* and *tp1031*) as the most suitable for MLST typing. These genes are proposed here as a new molecular typing scheme for *T. pallidum*. Each of these loci have been previously shown to contain recombinant regions, resulting from exchange among different *T. pallidum* subspecies [37,55–57]. These recombinant regions are very helpful in increasing the level of discrimination for the selected gene fragments included in the new MLST scheme. Incorporating these genes, which exhibit significant heterozygosity, will facilitate forthcoming genetic analyses based on MLST haplotypes.

The proposed *T. pallidum* MLST scheme shares six loci *(tp0136, tp0326, tp0548, tp0705, tp0858* and *tp0865*) with previous typing schemes available for the different subspecies of *T. pallidum* [1,12,15,16,31,32] (see Table 1). Hence, this novel MLST approach can be regarded as a refined and unified version of previous typing schemes which is now applicable to all subspecies of *T. pallidum*, after the incorporation of *tp1031*. It would be highly valuable to assess the resolution achieved by the new scheme proposed in this study compared to all previous typing schemes in future studies.

The efficiency of amplification in typing schemes is influenced by various factors, such as the type of specimen, bacterial load, time interval between sample collection and DNA isolation, DNA extraction technique, amplification protocol, and length of the amplification product. To enhance amplification efficiency, we opted for relatively shorter amplicons of 398-647 bp, compared to previous typing schemes. As a result, we were able to amplify samples using a single PCR instead of nested PCRs, which significantly reduced time and costs as well as improved efficiency. Through our approach, we were able to obtain full STs for 45% (82/183) of clinical samples analyzed in experimental laboratory settings. In contrast, we obtained a higher efficiency (82%, 151/183) for the amplification of the 23S rRNA gene compared to that obtained for the seven genes of the new MLST scheme. These rates are comparable to the amplification efficiency observed in previous *T. pallidum* typing studies, which ranged from 14 % to 95% with a median of 64.4% (Supplementary Table 16). However, a limitation of comparing these studies is that they employed different criteria for sample collection and investigation of *T. pallidum*, which could introduce heterogeneity in the efficiency obtained for each study.

It is important to note that a significant portion of the samples used in this study consisted of remnants from previous investigations, resulting in limited quantities of DNA with potentially compromised quality. These factors may have significantly impacted the efficiency achieved in our analyses, particularly for the seven genes included. In many cases, we were able to obtain the sequence of the 23S rRNA gene because it was the first one that was amplified, but not the sequence of some of the genes in the MLST scheme due to insufficient sample volume or poor DNA quality. As a result, the efficiency of obtaining the 23S rRNA gene sequence was higher compared to all the genes of the MLST scheme. Therefore, to obtain a more reliable assessment of efficiency, it is crucial to conduct further testing using recently collected samples.

We encountered some issues with the amplification efficiency of primers targeting the *tp0858* and *tp0865* genes in TEN samples, which led us to design new primers for these genes. The new primers for the *tp0858* gene were successfully tested on a set of TEN samples and we identified a deletion in the forward primer binding site that hindered amplification in these samples (Supplementary Figure 3). In contrast, the newly designed primers for the *tp0865* gene, while able to amplify a Nichols strain sample, did not amplify the target region in the TEN samples. To determine the effectiveness of the entire set of new and old primers for the *tp0865* gene, additional testing with more TEN samples and with a larger dataset of complete genomes is required. It is possible that in this set of TEN samples from Japan, there is an uncharacterized indel or SNP that disrupts primer binding, a hypothesis that might be resolved through the analysis of additional TEN genomes. In the meantime, our findings support the utility of the primers listed in Supplementary Table 6, unless samples exhibit amplification issues for the *tp0858* or *tp0865* genes. In such cases, it may be beneficial to use the primer combinations outlined in Supplementary Tables 8 and 9, respectively.

In our study, we applied the new typing scheme to analyze a total of 542 samples, including those subjected to typing through Sanger sequencing and those derived from WGS data. The results demonstrated a high level of discrimination, as we identified 82 different allelic profiles in 415 out of the 542 samples. However, we encountered challenges in assigning STs to 127 samples. These challenges arose primarily from unsuccessful gene amplification in the experimentally typed samples and a significant amount of missing data in the gene sequences from WGS samples. Our findings revealed that, among the 82 STs identified, 29 belonged to the SS14 clade, 31 to the Nichols clade, 16 to TPE, and 6 to TEN. Nevertheless, values of genetic diversity derived from the frequency distributions of the different STs within subspecies/sublineages did not show major differences (ranging from 0.92 to 0.75, Table 6), suggesting overall similar levels of genetic diversity. However, ST diversity in SS14-TPA was the lowest among them, likely reflecting the recent epidemic expansion of this lineage [19,37].

Consistent with prior research, our findings reveal a high proportion of macrolide-resistant strains (51% of samples) among those tested for resistance alleles (393 out of 542 samples) [13,19,51,58–61]. This proportion is far less than the 98% macrolide-resistant strains reported by Lieberman *et al*. [62], though specimens in the present study were collected over a broader geographic outside of North America and over a broader time span (1990’s-2020’s). As a result, first and second-line therapy for syphilis with azithromycin is no longer recommended as an alternative treatment to penicillin [63]. This underscores the necessity for further epidemiological investigations aimed at monitoring and characterizing the escalating dissemination of macrolide resistance in *T. pallidum*. Among the samples tested for resistance (393/542), a remarkable 84% of the resistant strains were identified as belonging to the SS14-sublineage (167/199). Additionally, resistance was detected in 12 Nichols-sublineage strains (12/126), 9/19 from TEN, and 11/19 from TPA strains with undefined sublineage assignment, whereas no resistance was observed in TPE. Most resistant samples (193) exhibited the A2058G mutation, whereas only 6 resistant samples (6/199) displayed the A2059G mutation. Among these six samples, five originated from the SS14-sublineage, and one from the Nichols-sublineage. Notably, we encountered challenges in determining the macrolide sensitivity or resistance of 117 samples from the analysis of WGS data, as well as for 32 samples evaluated by Sanger sequencing. This was mainly due to substantial missing data in the 23S rRNA gene sequences derived from WGS data and difficulties in amplifying this gene for some clinical samples, likely due to low DNA quality and/or quantity. Overall, these findings underscore the ongoing challenges posed by macrolide resistance in *T. pallidum* and emphasize the need for comprehensive epidemiological investigations and genomic studies to inform public health efforts and combat the rising spread of macrolide resistance.

The phylogenetic tree reconstructed using the concatenated MLST data is congruent with the differentiation of the clades and subspecies of *T. pallidum* (Figure 2). However, a comprehensive phylogeny of whole genomes that encompasses all samples typed by the new MLST scheme in this project is currently unavailable and such phylogeny would be necessary for a thorough comparison of the resolution provided by MLST versus whole genomes. Nonetheless, considering the number of distinct sequence types (82) obtained, it can be hypothesized that the resolution offered by MLST is quite high. When comparing the genetic variation obtained from the other alternative typing schemes currently available for *T. pallidum* (Table 1), it is essential to note that they vary not only in the number of loci utilized but also in their target regions and even the *T. pallidum* subspecies they target. Therefore, a direct comparison of the number of distinct STs identified between these schemes and the new MLST scheme proposed in this study is not appropriate.

Interestingly, in this study we included 11 samples that were sequenced more than once, using different methods or in different laboratories, to assess the consistency of the typing scheme (see Supplementary Note 1 for more details). All replicates were assigned to the same sequence type (ST), except for sample CDC2. This discrepancy was due to a single allelic difference in the *tp0136* gene, likely caused by variation within a microsatellite region. These findings support the overall robustness of the typing scheme, while also pointing to rare cases of variability that may result from biological or technical factors.

We observed genetic variation both within and between populations based on geographic origin, highlighting a degree of population structuring in *T. pallidum*. As expected, overall nucleotide diversity between continents was higher than within continents, suggesting geographic differentiation. The highest within-group variation was observed in the Americas, whereas Europe, despite including the largest and most geographically diverse sample set, showed the lowest within-continent diversity. At the country level, a similar trend was observed: Italy and Cuba displayed the highest within-country nucleotide diversity, while Ireland and Portugal showed the lowest. Madagascar, although the country with the largest number of samples, exhibited moderate diversity, pointing to the predominance of a few dominant lineages rather than broad genetic heterogeneity. The most frequent sequence types (STs) also reflected this regional pattern: ST20 was predominant in the USA, and ST57 in Madagascar, suggesting localized expansions.

Together, these results support the existence of clear geographic clustering in *T. pallidum*, with varying degrees of diversity shaped by regional transmission dynamics, sampling density, and historical patterns. Nonetheless, interpretation should remain cautious due to disparities in sample size, geographic distribution, and time of collection across regions.

Beale and coworkers [19] suggested weak overall geographic structuring for TPA, but they also noted that genetic diversity among *T. pallidum* populations is nonetheless influenced by geographic factors, indicating distinct population structures. They found frequent sharing of sublineages and genetically similar strains in countries with extensive sampling, suggesting that similar patterns might be found globally. It is possible, nonetheless, that population genetic structure is somewhat masked in phylogenetic analyses of vertically transmitted genes: in such analyses, recombinant regions are excluded. However, these regions are often very variable, and the exchanges can increase or decrease the similarities of strains, influencing genetic structure.

In addition to the geographic structuring of diversity, Beale et *al*. [19] also observed that some sublineages may be rare while others are more common, possibly due to fitness advantages or specific transmission dynamics, leading to private sublineages reflecting localized transmission networks. These results underscore complex transmission dynamics influenced by geographic and epidemiological factors, highlighting the importance of comprehensive sampling strategies. Our observations align with Beale *et al*.’s findings, indicating divergence of *T. pallidum* strains over time in different geographic regions, emphasizing the need to consider regional factors and transmission dynamics when analyzing pathogen spread. Further research is warranted to investigate transmission routes, conduct comparative analyses, and perform longitudinal studies to enhance our understanding of *T. pallidum* dynamics and inform targeted control measures.

In summary, we have developed a novel multi-locus typing (MLST) scheme utilizing genome-wide data from 121 *T. pallidum* genomes to scan for optimal typing loci. This scheme offers improved resolution for differentiating between *T. pallidum* subspecies and within TPA lineages, based on sequencing seven loci and analyzing 23S rRNA genes for macrolide resistance/sensitivity. Importantly, all the target regions can be amplified simultaneously using a simple PCR protocol, making it suitable even for scarce and low-quality samples. The scheme is publicly accessible in the PubMLST database [50], encouraging widespread adoption in standard laboratories due to its ease and speed of implementation. We believe this tool could advance *T. pallidum* epidemiology, facilitating longitudinal studies, infection tracking, and association identification with patient groups. These advancements will enhance public health interventions and establish a global molecular surveillance network with publicly accessible data [18].

## Supporting information

Suppementary_Figures

## ACKNOWLEDGMENTS

MPD has been funded by program FPU17/02367 from the Spanish Ministerio de Educación. This research has been supported by grants BFU2017-89594-R and PID2021-127010OB-I00 from Spanish MICIN, CIPROM2021-053 from Generalitat Valenciana (to FGC), by the Swiss National Science Foundation: grant number 188963 - “Towards the origins of syphilis” (to VJS), and by the University of Zurich’s University Research Priority Program “Evolution in Action: From Genomes to Ecosystems” (to VJS). Work in the Giacani’s lab was supported by the European Research Council (ERC) under the European Union’s Horizon 2020 Research and Innovation Program (Grant agreement No. 850450 to OM), NIH U19 AI144133 and by the Open Philanthropy Pledge #8394150 (to LG). Some experiments conducted in Japan were supported by Grants-in-Aid for research on HIV/AIDS from the Ministry of Health, Labor and Welfare of Japan (grant number 20HB1003) to TK. The work in the Czech Republic was partially funded by the National Institute of Virology and Bacteriology project (Programme EXCELES, ID Project No. LX22NPO5103, Funded by the European Union - Next Generation EU, recipient D.Š.).

## REFERENCES

1. Noda AA, Grillová L, Lienhard R, Blanco O, Rodríguez I, Šmajs D. Bejel in Cuba: molecular identification of Treponema pallidum subsp. endemicum in patients diagnosed with venereal syphilis. Clin Microbiol Infect. 2018;24: 1210.e1–1210.e5.

2. Shinohara K, Furubayashi K, Kojima Y, Mori H, Komano J, Kawahata T. Clinical perspectives of Treponema pallidum subsp. endemicum infection in adults, particularly men who have sex with men in the Kansai area, Japan: A case series. J Infect Chemother. 2022;28: 444–450.

3. Grange PA, Allix-Beguec C, Chanal J, Benhaddou N, Gerhardt P, Morini J-P, et al. Molecular subtyping of Treponema pallidum in Paris, France. Sex Transm Dis. 2013;40: 641–644.

4. World Health Organization. Global progress report on HIV, viral hepatitis and sexually transmitted infections, 2021. 2021.

5. Health Organization W. Summary report of a consultation on the eradication of yaws, 5-7 March 2012, Morges, Switzerland. 2012. Available: https://apps.who.int/iris/bitstream/handle/10665/75528/WHO?sequence=1

6. Mitjà O, Godornes C, Houinei W, Kapa A, Paru R, Abel H, et al. Re-emergence of yaws after single mass azithromycin treatment followed by targeted treatment: a longitudinal study. Lancet. 2018;391: 1599–1607.

7. Dofitas BL, Kalim SP, Toledo CB, Richardus JH. Yaws in the Philippines: first reported cases since the 1970s. Infect Dis Poverty. 2020;9: 1.

8. Elo A, Dégboé B, Barogui Y, Gomido IC, Wadagni A, d’Almeida C, et al. Resurgence of yaws in Benin: Four confirmed cases in the district of Z, Southern Benin. Journal of public health and epidemiology. 2019;11: 201–208.

9. Timothy JWS, Beale MA, Rogers E, Zaizay Z, Halliday KE, Mulbah T, et al. Epidemiologic and Genomic Reidentification of Yaws, Liberia. Emerg Infect Dis. 2021;27: 1123–1132.

10. Mitjà O, Šmajs D, Bassat Q. Advances in the diagnosis of endemic treponematoses: yaws, bejel, and pinta. PLoS Negl Trop Dis. 2013;7: e2283.

11. Pillay A, Liu H, Chen CY, Holloway B, Sturm WA, Steiner B, et al. Molecular Subtyping of Treponema pallidum Subspecies pallidum. Sexually Transmitted Diseases. 1998. pp. 408–414. doi:10.1097/00007435-199809000-00004

12. Grillová L, Bawa T, Mikalová L, Gayet-Ageron A, Nieselt K, Strouhal M, et al. Molecular characterization of Treponema pallidum subsp. pallidum in Switzerland and France with a new multilocus sequence typing scheme. PLoS One. 2018;13: e0200773.

13. Lieberman NAP, Lin MJ, Xie H, Shrestha L, Nguyen T, Huang M-L, et al. Treponema pallidum genome sequencing from six continents reveals variability in vaccine candidate genes and dominance of Nichols clade strains in Madagascar. PLoS Negl Trop Dis. 2021;15: e0010063.

14. Vrbová E, Pospíšilová P, Dastychová E, Kojanová M, Kreidlová M, Rob F, et al. Majority of Treponema pallidum ssp. pallidum MLST allelic profiles in the Czech Republic (2004-2022) belong to two SS14-like clusters. Sci Rep. 2024;14: 17463.

15. Marra CM, Sahi SK, Tantalo LC, Godornes C, Reid T, Behets F, et al. Enhanced molecular typing of treponema pallidum: geographical distribution of strain types and association with neurosyphilis. J Infect Dis. 2010;202: 1380–1388.

16. Godornes C, Giacani L, Barry AE, Mitja O, Lukehart SA. Development of a Multilocus Sequence Typing (MLST) scheme for Treponema pallidum subsp. pertenue: Application to yaws in Lihir Island, Papua New Guinea. Norris SJ, editor. PLoS Negl Trop Dis. 2017;11: e0006113.

17. Katz SS, Chi K-H, Nachamkin E, Danavall D, Taleo F, Kool JL, et al. Molecular strain typing of the yaws pathogen, Treponema pallidum subspecies pertenue. Kalendar R, editor. PLoS One. 2018;13: e0203632.

18. Thurlow CM, Joseph SJ, Ganova-Raeva L, Katz SS, Pereira L, Chen C, et al. Selective Whole-Genome Amplification as a Tool to Enrich Specimens with Low Treponema pallidum Genomic DNA Copies for Whole-Genome Sequencing. mSphere. 2022;7: e0000922.

19. Beale MA, Marks M, Cole MJ, Lee M-K, Pitt R, Ruis C, et al. Global phylogeny of Treponema pallidum lineages reveals recent expansion and spread of contemporary syphilis. Nat Microbiol. 2021;6: 1549–1560.

20. Edmondson DG, Delay BD, Kowis LE, Norris SJ. Parameters affecting continuous in vitro culture of Treponema pallidum strains. MBio. 2021;12: 1–21.

21. Pereira LE, Katz SS, Sun Y, Mills P, Taylor W, Atkins P, et al. Successful isolation of Treponema pallidum strains from patients’ cryopreserved ulcer exudate using the rabbit model. PLoS One. 2020;15: e0227769.

22. Maiden MC, Bygraves JA, Feil E, Morelli G, Russell JE, Urwin R, et al. Multilocus sequence typing: a portable approach to the identification of clones within populations of pathogenic microorganisms. Proc Natl Acad Sci U S A. 1998;95: 3140–3145.

23. Enright MC, Day NP, Davies CE, Peacock SJ, Spratt BG. Multilocus sequence typing for characterization of methicillin-resistant and methicillin-susceptible clones of Staphylococcus aureus. J Clin Microbiol. 2000;38: 1008–1015.

24. Dingle KE, Colles FM, Wareing DR, Ure R, Fox AJ, Bolton FE, et al. Multilocus sequence typing system for Campylobacter jejuni. J Clin Microbiol. 2001;39: 14–23.

25. Enright MC, Spratt BG. A multilocus sequence typing scheme for Streptococcus pneumoniae: identification of clones associated with serious invasive disease. Microbiology. 1998;144 (Pt 11): 3049–3060.

26. Katz KA, Pillay A, Ahrens K, Kohn RP, Hermanstyne K, Bernstein KT, et al. Molecular Epidemiology of Syphilis—San Francisco, 2004-2007. Sexually Transmitted Diseases. 2010. pp. 660–663. doi:10.1097/olq.0b013e3181e1a77a

27. Mikalová L, Pospíšilová P, Woznicová V, Kuklová I, Zákoucká H, Smajs D. Comparison of CDC and sequence-based molecular typing of syphilis treponemes: tpr and arp loci are variable in multiple samples from the same patient. BMC Microbiol. 2013;13: 178.

28. Grillova L, Jolley K, Šmajs D, Picardeau M. A public database for the new MLST scheme for Treponema pallidum subsp. pallidum : surveillance and epidemiology of the causative agent of syphilis. PeerJ. 2019;6: e6182.

29. Flasarová M, Smajs D, Matejková P, Woznicová V, Heroldová-Dvoráková M, Votava M. [Molecular detection and subtyping of Treponema pallidum subsp. pallidum in clinical specimens]. Epidemiol Mikrobiol Imunol. 2006;55: 105–111.

30. Woznicová V, Smajs D, Wechsler D, Matĕjková P, Flasarová M. Detection of Treponema pallidum subsp. pallidum from skin lesions, serum, and cerebrospinal fluid in an infant with congenital syphilis after clindamycin treatment of the mother during pregnancy. J Clin Microbiol. 2007;45: 659–661.

31. Chuma IS, Roos C, Atickem A, Bohm T, Anthony Collins D, Grillová L, et al. Strain diversity of Treponema pallidum subsp. pertenue suggests rare interspecies transmission in African nonhuman primates. Sci Rep. 2019;9: 14243.

32. Medappa M, Pospíšilová P, Madruga MPM, John LN, Beiras CG, Grillová L, et al. Low genetic diversity of Treponema pallidum ssp. pertenue (TPE) isolated from patients’ ulcers in Namatanai District of Papua New Guinea: Local human population is infected by three TPE genotypes. PLoS Negl Trop Dis. 2024;18: e0011831.

33. Peltzer A, Jäger G, Herbig A, Seitz A, Kniep C, Krause J, et al. EAGER: efficient ancient genome reconstruction. Genome Biol. 2016;17: 60.

34. Li C. A Burrows-Wheeler Transform Based Method for DNA Sequence Comparison. Computational Biology and Bioinformatics. 2014. p. 33. doi:10.11648/j.cbb.20140203.11

35. Okonechnikov K, Conesa A, García-Alcalde F. Qualimap 2: advanced multi-sample quality control for high-throughput sequencing data. Bioinformatics. 2016;32: 292–294.

36. McKenna A, Hanna M, Banks E, Sivachenko A, Cibulskis K, Kernytsky A, et al. The Genome Analysis Toolkit: a MapReduce framework for analyzing next-generation DNA sequencing data. Genome Res. 2010;20: 1297–1303.

37. Arora N, Schuenemann VJ, Jäger G, Peltzer A, Seitz A, Herbig A, et al. Origin of modern syphilis and emergence of a pandemic Treponema pallidum cluster. Nat Microbiol. 2016;2: 16245.

38. Minh BQ, Schmidt HA, Chernomor O, Schrempf D, Woodhams MD, von Haeseler A, et al. IQ-TREE 2: New Models and Efficient Methods for Phylogenetic Inference in the Genomic Era. Mol Biol Evol. 2020;37: 1530–1534.

39. Strimmer K, von Haeseler A. Likelihood-mapping: a simple method to visualize phylogenetic content of a sequence alignment. Proc Natl Acad Sci U S A. 1997;94: 6815–6819.

40. Lukehart SA, Godornes C, Molini BJ, Sonnett P, Hopkins S, Mulcahy F, et al. Macrolide resistance in Treponema pallidum in the United States and Ireland. N Engl J Med. 2004;351: 154–158.

41. Staden R, Judge DP, Bonfield JK. Managing Sequencing Projects in the GAP4 Environment. Introduction to Bioinformatics. pp. 327–344. doi:10.1385/1-59259-335-6:327

42. Larsson A. AliView: a fast and lightweight alignment viewer and editor for large datasets. Bioinformatics. 2014;30: 3276–3278.

43. Matejkova P, Flasarova M, Zakoucka H, Borek M, Kremenova S, Arenberger P, et al. Macrolide treatment failure in a case of secondary syphilis: a novel A2059G mutation in the 23S rRNA gene of Treponema pallidum subsp. pallidum. J Med Microbiol. 2009;58: 832–836.

44. Molini BJ, Tantalo LC, Sahi SK, Rodriguez VI, Brandt SL, Fernandez MC, et al. Macrolide Resistance in Treponema pallidum Correlates With 23S rDNA Mutations in Recently Isolated Clinical Strains. Sex Transm Dis. 2016;43: 579–583.

45. Katoh K, Asimenos G, Toh H. Multiple alignment of DNA sequences with MAFFT. Methods Mol Biol. 2009;537: 39–64.

46. Fu L, Niu B, Zhu Z, Wu S, Li W. CD-HIT: accelerated for clustering the next-generation sequencing data. Bioinformatics. 2012;28: 3150–3152.

47. Pla-Díaz M, Akgül G, Molak M, du Plessis L, Panagiotopoulou H, Doan K, et al. Insights into Treponema pallidum genomics from modern and ancient genomes using a novel mapping strategy. BMC Biol. 2025;23: 7.

48. Nei M. F-statistics and analysis of gene diversity in subdivided populations. Ann Hum Genet. 1977;41: 225–233.

49. Rozas J, Ferrer-Mata A, Sánchez-DelBarrio JC, Guirao-Rico S, Librado P, Ramos-Onsins SE, et al. DnaSP 6: DNA Sequence Polymorphism Analysis of Large Data Sets. Mol Biol Evol. 2017;34: 3299–3302.

50. Jolley KA, Bray JE, Maiden MCJ. Open-access bacterial population genomics: BIGSdb software, the PubMLST.org website and their applications. Wellcome Open Res. 2018;3: 124.

51. Taouk ML, Taiaroa G, Pasricha S, Herman S, Chow EPF, Azzatto F, et al. Characterisation of Treponema pallidum lineages within the contemporary syphilis outbreak in Australia: a genomic epidemiological analysis. The Lancet Microbe. 2022. pp. e417–e426. doi:10.1016/s2666-5247(22)00035-0

52. Vrbová E, Noda AA, Grillová L, Rodríguez I, Forsyth A, Oppelt J, et al. Whole genome sequences of Treponema pallidum subsp. endemicum isolated from Cuban patients: The non-clonal character of isolates suggests a persistent human infection rather than a single outbreak. PLoS Negl Trop Dis. 2022;16: e0009900.

53. Seña AC, Matoga MM, Yang L, Lopez-Medina E, Aghakhanian F, Chen JS, et al. Clinical and genomic diversity of Treponema pallidum subspecies pallidum to inform vaccine research: an international, molecular epidemiology study. Lancet Microbe. 2024;5: 100871.

54. Beale MA, Thorn L, Cole MJ, Pitt R, Charles H, Ewens M, et al. Genomic epidemiology of syphilis in England: a population-based study. Lancet Microbe. 2023;4: e770–e780.

55. Pla-Díaz M, Sánchez-Busó L, Giacani L, Šmajs D, Bosshard PP, Bagheri HC, et al. Evolutionary Processes in the Emergence and Recent Spread of the Syphilis Agent, Treponema pallidum. Mol Biol Evol. 2022;39. doi:10.1093/molbev/msab318

56. Grillová L, Oppelt J, Mikalová L, Nováková M, Giacani L, Niesnerová A, et al. Directly Sequenced Genomes of Contemporary Strains of Syphilis Reveal Recombination-Driven Diversity in Genes Encoding Predicted Surface-Exposed Antigens. Front Microbiol. 2019;10: 1691.

57. Strouhal M, Mikalová L, Haviernik J, Knauf S, Bruisten S, Noordhoek GT, et al. Complete genome sequences of two strains of Treponema pallidum subsp. pertenue from Indonesia: Modular structure of several treponemal genes. PLoS Negl Trop Dis. 2018;12: e0006867.

58. Beale MA, Marks M, Sahi SK, Tantalo LC, Nori AV, French P, et al. Genomic epidemiology of syphilis reveals independent emergence of macrolide resistance across multiple circulating lineages. Nat Commun. 2019;10: 1–9.

59. Obraztsova O, Shpilevaya MV, Katunin G, Obukhov A, Shagabieva YZ, Solomka V. Prevalence of the A2058G mutation in 23S rRNA gene, which determines Treponema pallidum macrolide resistance in Russian population. Clin Microbiol Antimicrob Chemother. 2022. doi:10.36488/cmac.2022.4.369-374

60. Morando N, Vrbová E, Melgar A, Rabinovich RD, Šmajs D, Pando MA. High frequency of Nichols-like strains and increased levels of macrolide resistance in Treponema pallidum in clinical samples from Buenos Aires, Argentina. Scientific Reports. 2022. doi:10.1038/s41598-022-20410-5

61. Wang X, Abliz P, Deng S. Molecular Characteristics of Macrolide Resistance in from Patients with Latent Syphilis in Xinjiang, China. Infect Drug Resist. 2023;16: 1231–1236.

62. Lieberman NAP, Reid TB, Cannon CA, Nunley BE, Berzkalns A, Cohen SE, et al. Near-Universal Resistance to Macrolides of Treponema pallidum in North America. N Engl J Med. 2024;390: 2127–2128.

63. Janier M, Hegyi V, Dupin N, Unemo M, Tiplica GS, Potočnik M, et al. 2014 European guideline on the management of syphilis. J Eur Acad Dermatol Venereol. 2014;28: 1581–1593.

