## Supplementary_Material for "A new typing scheme demonstrates high discriminatory power for *Treponema pallidum* subspecies"

(1) Joint Research Unit Infection and Public Health FISABIO-Univ. Valencia, Institute for Integrative Systems Biology (I2SysBio, CSIC-UV),Valencia, Spain.
(2) Department of Medicine, Division of Allergy and Infectious Diseases, University of Washington, Seattle WA, USA.
(3) Department of Neurology, University of Washington School of Medicine, Seattle, WA, USA.
(4) Department of Biology, Faculty of Medicine, Masaryk University, Brno, Czech Republic.
(5) Division of Microbiology, Osaka Institute of Public Health, Osaka, Japan.
(6) Centers for Disease Control and Prevention, Atlanta, GA, USA.
(7) National Reference Laboratory of Treponemes and Special Pathogens, Tropical Medicine Institute "Pedro Kourí", Havana, Cuba.
(8) Dermatology Clinic, University Hospital Zurich, Zurich, Switzerland.
(9) School of Translational Medicine, Monash University, Melbourne, Victoria, Australia.
(10) ISGlobal, Barcelona, Spain.
(11) Institute of Evolutionary Medicine, University of Zurich, Zurich, Switzerland.
(12) Department of Environmental Sciences, University of Basel, Basel, Switzerland.
(13) Institute for Bioinformatics and Medical Informatics, University of Tübingen, Tübingen, Germany.
(14) Dermatology Service, Consorcio Hospital General Universitario, Valencia, Spain.
(15) CEU Cardenal Herrera University of Castellón, Castellón, Spain.
(16) Microbiology Service, Consorcio Hospital General Universitario, Valencia, Spain.
(17) Catholic University of Valencia San Vicente Mártir, Valencia, Spain.
(18) Zurich Institute of Forensic Medicine, University of Zurich, Zurich, Switzerland.
(19) CIBER in Epidemiology and Public Health, Madrid, Spain.

### * Correspondence to:

### INDEX OF CONTENT

### SUPPLEMENTARY NOTES

#### Supplementary Note 1

In this study, we included duplicate sequences from 10 samples that were resequenced. These sequences were obtained from the same DNA isolate but sequenced using different methods or in a different laboratory (Supplementary Note 1, Table 1). These duplicates were analyzed to identify any sequence differences, potentially arising during the laboratory and bioinformatic workflows, that might result in different sequence types (STs) despite their common origin. The genome sequences of samples Sea81_4_1 and Sea81_4_2, obtained through resequencing the original DNA sample isolated in 1951 [1] and subjected to different culture passages in rabbits, are still unpublished (Supplementary Note 1, Table 1).

**Supplementary Note 1, Table 1.** Details for the duplicate sequences from the 10 samples that were resequenced. A comparison of the allelic profiles and the ST obtained for each resequenced sample (rows) is shown in the table. The allelic profile for each gene is provided in the following order: *tp0136*, *tp0326*, *tp0548*, *tp0705*, *tp0858*, *tp0865* and *tp1031* and is specified for each sample. Differences observed in the allelic profiles are highlighted in red. Samples for which the sequence data was obtained through Sanger sequencing of clinical samples are indicated in blue; samples for which the data was obtained from the original assembly file obtained by *de novo* assembly are indicated in green. The remaining samples are highlighted in orange because their gene sequences were obtained through processing of reads obtained in the different studies, as referenced, by mapping using Nichols as genome reference.

| Sequence 1 | Raw reads source | Allelic profile | ST | Sequence 2 | Raw reads source | Allelic profile | ST |
|---|---|---|---|---|---|---|---|
| **C279_1** | [2] | 3,14,34,4,3,5,4 | 67 | **C279_2** | - | 3,14,34,4,3,5,4 | 67 |
| **CDC2** | [3] | **16**,6,6,2,4,4,4 | 10 | **CDC2_2** | - | **4,**,6,6,2,4,4,4 | 82 |
| **SS14_1** | [4] | 1,1,12,2,1,1,1 | 20 | **SS14_2** | - | 1,1,12,2,1,1,1 | 20 |
| **GHA1** | [5] | 16,15,23,2,15,4,4 | 12 | **Ghana_051** | [6] | 16,15,23,2,15,4,4 | 12 |
| **CDC-1** | [5] | 16,15,23,2,15,4,4 | 12 | **CDC2575** | [6] | 16,15,23,2,15,4,4 | 12 |
| **GRA2** | [5] | 1,1,12,2,1,1,1 | 20 | **Grady** | [7] | 1,1,12,2,1,1,1 | 11 |
| **IND1** | [5] | 16,25,39,7,27,18,4 | 44 | **Kampung_Dalan_363** | [8] | 16,25,39,7,27,18,4 | 44 |
| **Sea81_4_1** | Unpublished | 5,18,22,2,20,13,2 | 16 | **Seattle81** | [1,7] | 5,18,22,2,20,13,2 | 16 |
| **Sea81_4_2** | Unpublished | 5,18,22,2,20,13,2 | 16 | | | | |
| **NIC1** | [5] | 2,2,2,2,2,2,2 | 2 | **Nichols_1**/ **Nichols_2*** | [4]/ this study | 2,2,2,2,2,2,2 | 2 |
| **NIC2** | [5] | 2,2,2,2,2,2,2 | 2 | | | | |
| **SAM1** | [5] | 4,16,8,2,25,9,4 | 9 | SAMD | [3] | 4,16,8,2,25,9,4 | 9 |

As shown in Supplementary Note 1, Table 1, all samples, except for the CDC2 replicates, were assigned to the same ST. This sample was sequenced in two different studies. In one, it was obtained via de novo assembly after the original material had been passaged in rabbits [3]. In the other, analyzed in the present study, one of the aliquots from the original DNA sample was sequenced using Sanger technology.

Despite originating from the same biological material, the two aliquots produced different STs due to a single allelic difference in the *tp0136* gene: one carried allele 4, the other allele 16. Upon examination, this discrepancy was found to result from three SNPs located within a stretch of repeated CTT motifs.

We suspect this region corresponds to a microsatellite, which may be prone to instability. Such variability could result from replication slippage during bacterial growth, possibly introduced during rabbit passage, or reflect intra-sample heterogeneity captured differently depending on the sequencing approach. Alternatively, the observed difference might stem from technical artifacts during library preparation or assembly, as repetitive regions like microsatellites are often difficult to resolve accurately.

### SUPPLEMENTARY FILES

**Supplementary File 1.** In-house script to compute the number of SNPs per gene.

**Supplementary File 2.** In-house script used to determine the subspecies and/or clades distinguishable by each of these SNPs.

**Supplementary File 3.** In-house script used to discard haplotypes that could not be considered due to missing data in the sequences employed.

**Supplementary File 4.** The number of different alleles obtained for the gene *tp0136*

**Supplementary File 5.** The number of different alleles obtained for the gene *tp0326*

**Supplementary File 6.** The number of different alleles obtained for the gene *tp0548*

**Supplementary File 7.** The number of different alleles obtained for the gene *tp0705*

**Supplementary File 8.** The number of different alleles obtained for the gene *tp0858*

**Supplementary File 9.** The number of different alleles obtained for the gene *tp0865*

**Supplementary File 10.** The number of different alleles obtained for the gene *tp1031*

### SUPPLEMENTARY TABLES

**Supplementary Table 1A.** The 121 *in silico* genomes employed for the design of the new MLST scheme for *T. pallidum*.

**Supplementary Table 1B.** Supplementary Table 1B. The 238 publicly available in silico genome assemblies used for the *in silico* application of the newly designed MLST scheme.

**Supplementary Table 2.** Complete results of the likelihood mapping test for the MSA using Nichols as reference genome for mapping. Of the 978 protein-coding genes, 332 showed some phylogenetic signal and were retained for the ensuing analyses. It was not possible to perform this test for 198 genes due to the large number of undetermined positions in the corresponding multiple alignments. (Zones 1-3 represent cases in which one topology has a significantly higher likelihood than the other two alternative topologies; zones 4-6 represent cases in which one topology has a significantly lower likelihood than the other two, and zone 7 represents cases in which all the topologies have similar likelihoods, hence the corresponding gene does not carry enough phylogenetic signal to differentiate between the 3 evolutionary hypotheses tested in each case).

**Supplementary Table 3**. Number of SNPs obtained per each gene selected in the likelihood mapping test.

**Supplementary Table 4.** The results of the in-house script employed to know which subspecies and/or clades could be differentiated by each SNPs per gene. The excel file includes 21 different sheets, 20 corresponding the results of this script for the final candidate genes for primers design, plus the additional gene *tp0705*. The first column of each sheet represents the variable positions for that gene, and the following columns show which subspecies or clades it is possible to differentiate according to which SNPs and which strains have SNPs at that position. A summary graph of the results of that script by gene is also shown.

**Supplementary Table 5.** Minimum and theoretical number of haplotypes (ST) to differentiate the final 16 genes selected for the new MLST scheme. Note that some genes have more than one possible primer combination and all of them have been analyzed and included in this table. When the gene has more than one possible primer combination, it is indicated by different numbers. For each sample and locus, the different alleles determined are indicated in different colors, as well as the final haplotype (ST) corresponding to each allelic combination for the 16 loci. This table does not include the 23S gene. Alleles that could not be assigned due to the presence of Ns in the sequences of those genes for those samples are highlighted in black. Please note that both the allele and ST assignments shown in this table are provisional and were used solely for the design and evaluation of the MLST scheme. They are intended for illustrative purposes and do not correspond to the final validated allele definitions or ST assignments.

**Supplementary Table 6.** Final primers designed for the new *T. pallidum* MLST scheme. The primers for the *23S* rRNA gene are the same primers used in [9] to complete the typing MLST scheme and to know if the samples are resistant to tetracycline or not. The amplicon size is the final size obtained after Sanger sequencing.

| Gene | Primer sequence | Melting Temperature | Amplicon Size (bp) |
|---|---|---|---|
| ***tp0136*** | 5'- AGCGACGGGTGCTATCACTA -3' | 58.75ºC | 566 |
| | 5'- TTACTCGCGGTTCCAGGAGC -3' | | |
| ***tp0326*** | 5'- CATTCGTTTCGCTCCGACAC -3' | 55.5ºC | 545 |
| | 5'- TACCGTGAACGACAACACAA -3' | | |
| ***tp0548*** | 5'- ATGATATCGTGTTCGGTGCG -3' | 55.8ºC | 506 |
| | 5'- ACAGAAGGTGTGAGACGCAT -3' | | |
| ***tp0705*** | 5'- ACCGACCATATCCAGTACAC -3' | 57.85ºC | 545 |
| | 5'- TCTTCTCTCACACACGTTGC -3' | | |
| ***tp0858*** | 5'- AAGTGTGGTTGCTGCAAGGA -3' | 57.55ºC | 452 |
| | 5'- ATTTCGGCCGAGCAGTATCG -3' | | |
| ***tp0865*** | 5'- GGCAATCGCTTCCTCATAGT -3' | 59ºC | 647 |
| | 5'- GGCATCAGTGTGGGAACCAA -3' | | |
| ***tp1031*** | 5'- TTGCTGAGCATGCAGTGGAA -3' | 57.85ºC | 398 |
| | 5'- CACGTGGTACTGCATTGCCT -3' | | |
| ***23S*** | 5'- GTACCGCAAACCGACACAG -3' | 59ºC | 592 |
| | 5'- AGTCAAACCGCCCACCTAC -3' | | |

**Supplementary Table 7.** The minimum and theoretical number of haplotypes (STs) that allows differentiating the 7 final genes selected for the new MLST scheme. For each sample and locus, the different alleles determined are indicated in different colors as well as the final haplotype (ST) that corresponds to each allele combination for the 7 loci. This table does not include the 23S gene. Alleles that could not be assigned due to the presence of Ns in the sequences of those genes for those samples are highlighted in black. The haplotypes (STs) indicated with (?) are STs not properly determined due the indetermination in some of their alleles (NA) because of Ns in that gene's sequences. Please note that both the allele and ST assignments shown in this table are provisional and were used solely for the design and evaluation of the MLST scheme. They are intended for illustrative purposes and do not correspond to the final validated allele definitions or ST assignments.

**Supplementary Table 8.** New alternative primers designed for loci *tp0858* and *tp0865*.

| Gene | Primer | Primer sequence | Melting Temperature | Amplicon size |
|---|---|---|---|---|
| ***tp0858*** | F_2 | 5’- ACCGTAAAGGTCTCGGACAA -3’ | 57.6 ºC | 544 pb |
| | R_2 | 5’- GTGCCCTGCTGAAGAATGCG -3’ | | |
| ***tp0865*** | F_2 | 5’- CACGCCCCGTATAAAGAACA -3’ | 55 ºC | - |
| | F_3 | 5’- GCAACCGCCGAGGGTGTCTT -3’ | 62.9 ºC | - |
| | R_2 | 5’- CCACCAGGAGATAGGGGAAC -3’ | 56.8 ºC | - |

**Supplementary Table 9.** Primer combinations used to test the new primers designed for the gene *tp0865* with the old ones.

| Gene | Combination | Primers | Primer sequence | Melting Temperature | Amplicon size |
|---|---|---|---|---|---|
| ***tp0865*** | 1 | F_2 | 5’- CACGCCCCGTATAAAGAACA -3’ | 56.5 ºC | 836 |
| | | R_1 | 5’- GGCATCAGTGTGGGAACCAA -3’ | | |
| | 2 | F_1 | 5’- GGCAATCGCTTCCTCATAGT -3’ | 55.9 ºC | 957 |
| | | R_2 | 5’- CCACCAGGAGATAGGGGAAC -3’ | | |
| | 3 | F_3 | 5’- GCAACCGCCGAGGGTGTCTT -3’ | 60.4 ºC | 903 |
| | | R_1 | 5’- GGCATCAGTGTGGGAACCAA -3’ | | |

**Supplementary Table 10.** Final allelic profiles obtained per gene obtained for each sample. The table also includes information on the country where the sample was collected, the source of the sample, and whether the allelic profile was determined through in-silico analysis or experimentally. The symbol "-" indicates those alleles that could not be determined for a given sample. The symbol # indicates that alleles 4 and 16 of the tp0136 gene differ by three SNPs within a repeated CTT motif, likely corresponding to a microsatellite. This variability may arise from replication slippage, rabbit passage, sequencing biases, or technical artifacts during assembly. Further confirmation is needed to clarify whether in silico samples carry allele 4 or 16. Alleles marked with an asterisk (*) contain unresolved bases; therefore, these alleles and their corresponding sequence types (STs) have not been submitted to pubMLST, as their sequences require future confirmation.

**Supplementary Table 11.** The STs obtained per each sample. The table also includes information on the country where the sample was collected, the source of the sample, and whether the allelic profile was determined through in-silico analysis or experimentally. It also specifies the subspecies or clade determined for each sample and the macrolide resistance profile determined. The symbol # indicates that alleles 4 and 16 of the *tp0136* gene differ by three SNPs within a repeated CTT motif, likely corresponding to a microsatellite. This variability may arise from replication slippage, rabbit passage, sequencing biases, or technical artifacts during assembly. Further confirmation is needed to clarify whether in silico samples carry allele 4 or 16. Alleles marked with an asterisk (*) contain unresolved bases; therefore, these alleles and their corresponding sequence types (STs) have not been submitted to pubMLST, as their sequences require future confirmation.

**Supplementary Table 12.** Average number of nucleotide differences (*k*) in a population and between populations at continental level. The table is coloured with the following codes: green for the lowest values, yellow for intermediate values and purple for the highest values. The number of samples (N) from each continent is also indicated.

| N | Continent | Europe | America | Africa | Asia |
|---|---|---|---|---|---|
| 124 | **Europe** | **5.29** | | | |
| 93 | **America** | 11.98 | **15.95** | | |
| 85 | **Africa** | 39.16 | 29.74 | **5.50** | |
| 79 | **Asia** | 8.19 | 12.85 | 39.40 | **6.22** |

**Supplementary Table 13.** Average number of nucleotide substitutions per site within the four continents considered (π), and number of nucleotide substitutions per site between continents with Jukes and Cantor correction, Da(JC) on the lower hemimatrix, and standard deviation of Da(JC) on the upper hemimatrix. The table is coloured with the following codes: green for the lowest values, yellow for intermediate values and purple for the highest values. The number of samples (N) from each continent is also indicated.

| N | Continent | Europe | America | Africa | Asia |
|---|---|---|---|---|---|
| 124 | **Europe** | **0.00185** | 0.00075 | 0.00122 | 0.00064 |
| 93 | **America** | 0.00048 | **0.00544** | 0.00111 | 0.00083 |
| 85 | **Africa** | 0.01023 | 0.007 | **0.00197** | 0.00145 |
| 79 | **Asia** | 0.00028 | 0.00053 | 0.00981 | **0.00210** |

**Supplementary Table 14.** Average number of nucleotide differences (k) in a population and between populations at country level. The table is coloured with the following codes: green for the lowest values, yellow for intermediate values and purple for the highest values.

| N | Country | Madagascar | USA | Japan | Spain | China | Portugal | Switzerland | Ireland | Cuba | Czechia | Italy |
|---|---|---|---|---|---|---|---|---|---|---|---|---|
| 85 | **Madagascar** | **9.43** | | | | | | | | | | |
| 69 | **USA** | 28.47 | **17.88** | | | | | | | | | |
| 53 | **Japan** | 39.54 | 14.19 | **7.88** | | | | | | | | |
| 40 | **Spain** | 40.90 | 13.21 | 6.66 | **3.29** | | | | | | | |
| 25 | **China** | 40.29 | 13.58 | 6.03 | 4.92 | **4.38** | | | | | | |
| 25 | **Portugal** | 41.13 | 12.26 | 5.27 | 1.99 | 3.44 | **0.52** | | | | | |
| 12 | **Switzerland** | 37.82 | 13.74 | 8.74 | 5.50 | 7.29 | 4.40 | **8.14** | | | | |
| 11 | **Ireland** | 41.52 | 12.53 | 5.59 | 1.76 | 3.80 | 0.97 | 4.14 | **0.18** | | | |
| 11 | **Cuba** | 36.14 | 16.17 | 14.54 | 11.76 | 13.28 | 10.85 | 13.11 | 10.60 | **19.67** | | |
| 10 | **Czechia** | 41.75 | 13.13 | 6.20 | 2.69 | 4.40 | 1.88 | 4.90 | 1.41 | 11.47 | **2.16** | |
| 10 | **Italy** | 36.45 | 16.56 | 15.63 | 13.19 | 14.52 | 12.29 | 14.20 | 12.14 | 19.22 | 13.00 | **21.64** |

**Supplementary Table 15.** Average number of nucleotide substitutions per site within the nine countries considered (π), and number of nucleotide substitutions per site between countries with Jukes and Cantor correction, Da(JC) on the lower hemimatrix, and standard deviation of Da(JC) on the upper hemimatrix, and standard deviation of Dxy(JC) on the upper hemimatrix. The table is coloured with the following codes: green for the lowest values, yellow for intermediate values and purple for the highest values.

| N | Country | Madagascar | USA | Japan | Spain | China | Portugal | Switzerland | Ireland | Cuba | Czechia | Italy |
|---|---|---|---|---|---|---|---|---|---|---|---|---|
| 85 | **Madagascar** | **0.00295** | 0.00118 | 0.00159 | 0.00148 | 0.00187 | 0.00161 | 0.00263 | 0.00228 | 0.0028 | 0.00208 | 0.00289 |
| 69 | **USA** | 0.00609 | **0.00577** | 0.00102 | 0.00096 | 0.00112 | 0.00099 | 0.00145 | 0.00124 | 0.00158 | 0.00107 | 0.00161 |
| 53 | **Japan** | 0.00983 | 0.00068 | **0.0022** | 0.00075 | 0.00094 | 0.00065 | 0.00148 | 0.00074 | 0.00235 | 0.00062 | 0.00258 |
| 40 | **Spain** | 0.0109 | 0.00091 | 0.00028 | **0.00092** | 0.00076 | 0.00041 | 0.00133 | 0.00046 | 0.00223 | 0.00038 | 0.00247 |
| 25 | **China** | 0.01055 | 0.00089 | -3E-05 | 0.0003 | **0.00122** | 0.00066 | 0.00151 | 0.00074 | 0.00241 | 0.00063 | 0.00267 |
| 25 | **Portugal** | 0.01136 | 0.00099 | 0.00028 | 0.00002 | 0.00027 | **0.00014** | 0.00131 | 0.00012 | 0.00226 | 0.00017 | 0.00254 |
| 12 | **Switzerland** | 0.00957 | 0.00051 | 0.00023 | -6E-05 | 0.0003 | 0.00002 | **0.00235** | 0.00147 | 0.00273 | -1E-05 | 0.00289 |
| 11 | **Ireland** | 0.01154 | 0.00113 | 0.00042 | 0 | 0.00042 | 0.00007 | 0 | **0.00005** | 0.00258 | 0.00017 | 0.0029 |
| 11 | **Cuba** | 0.00762 | 0.00008 | 0.00021 | 0.00008 | 0.00035 | 0.00021 | -0.00018 | 0.00019 | **0.0055** | 0.00198 | 0.00315 |
| 10 | **Czechia** | 0.01134 | 0.00104 | 0.00031 | -1E-05 | 0.00031 | 0.00003 | 0.00124 | -1E-05 | 0.00016 | **0.0006** | 0.00221 |
| 10 | **Italy** | 0.00686 | -0.0001 | 0.00023 | 0.00018 | 0.00042 | 0.00031 | -0.00015 | 0.00031 | -0.0004 | 0.00028 | **0.00605** |

**Supplementary Table 16.** Reported efficiency (%) of molecular typing across different previous studies, based on amplification or typing success rates in clinical samples.

| Efficiency (%) | Reference |
|---|---|
| 60 | [10] |
| 69.4 | [11] |
| 84.4 | [12] |
| 95.3 | [13] |
| 34.5 | [14] |
| 50 | [15] |
| 19 | [16] |
| 54 | [17] |
| 90 | [18] |
| 13.6 | [19] |
| 75 | [20] |
| 48 | [21] |
| 51 | [22] |
| 92.6 | [23] |
| 75 | [24] |
| 60 | [25] |
| 38.8 | [26] |
| 84.2 | [27] |
| 73.9 | [28] |
| 30 | [29] |
| 0 | [30] |
| 58 | [31] |
| 84.1 | [32] |
| 68.7 | [33] |
| 23.5 | [34] |
| 68.75 | [35] |
| 84.2 | [36] |
| 75 | [37] |

### SUPPLEMENTARY FIGURES

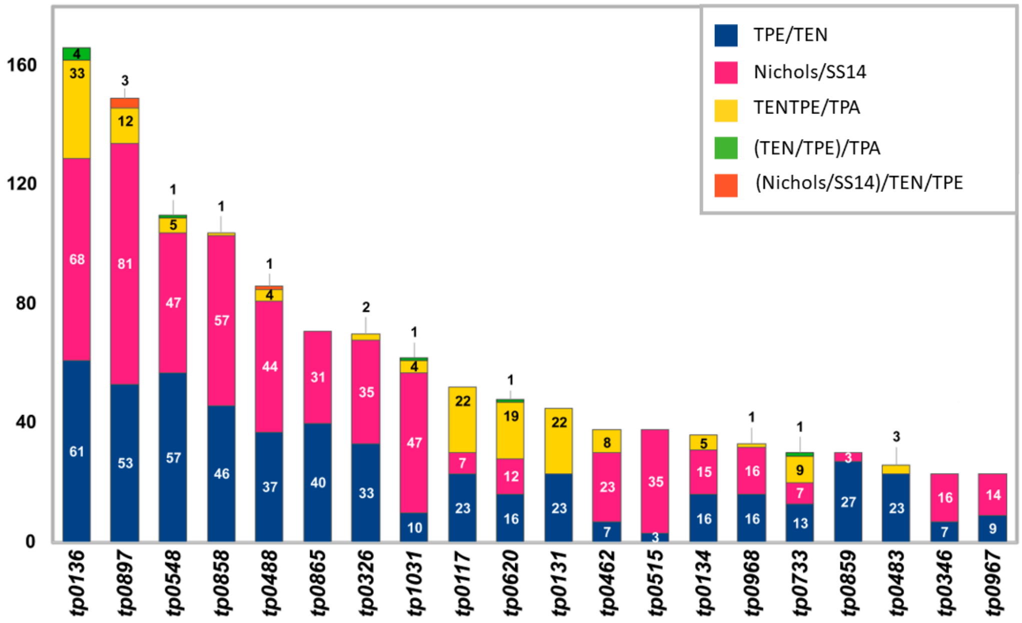

**Supplementary Figure 1.** Graph summarizing the discriminatory power at the different levels used to select 20 candidate genes for primer design. Numbers in each column sector represent the number of SNPs in each gene that can differentiate within or between subspecies/clades with the following code: dark blue, SNPs that differentiate between TPE and TEN; fuchsia, SNPs that differentiate Nichols and SS14; yellow, SNPs that differentiate TPE and TEN from TPA (but do not differentiate TPE from TEN); green, SNPs that differentiate TPE from TEN and in turn from TPA; and orange, SNPs that differentiate Nichols from SS14 and in turn from TPE and TEN (but do not differentiate TPE from TEN).

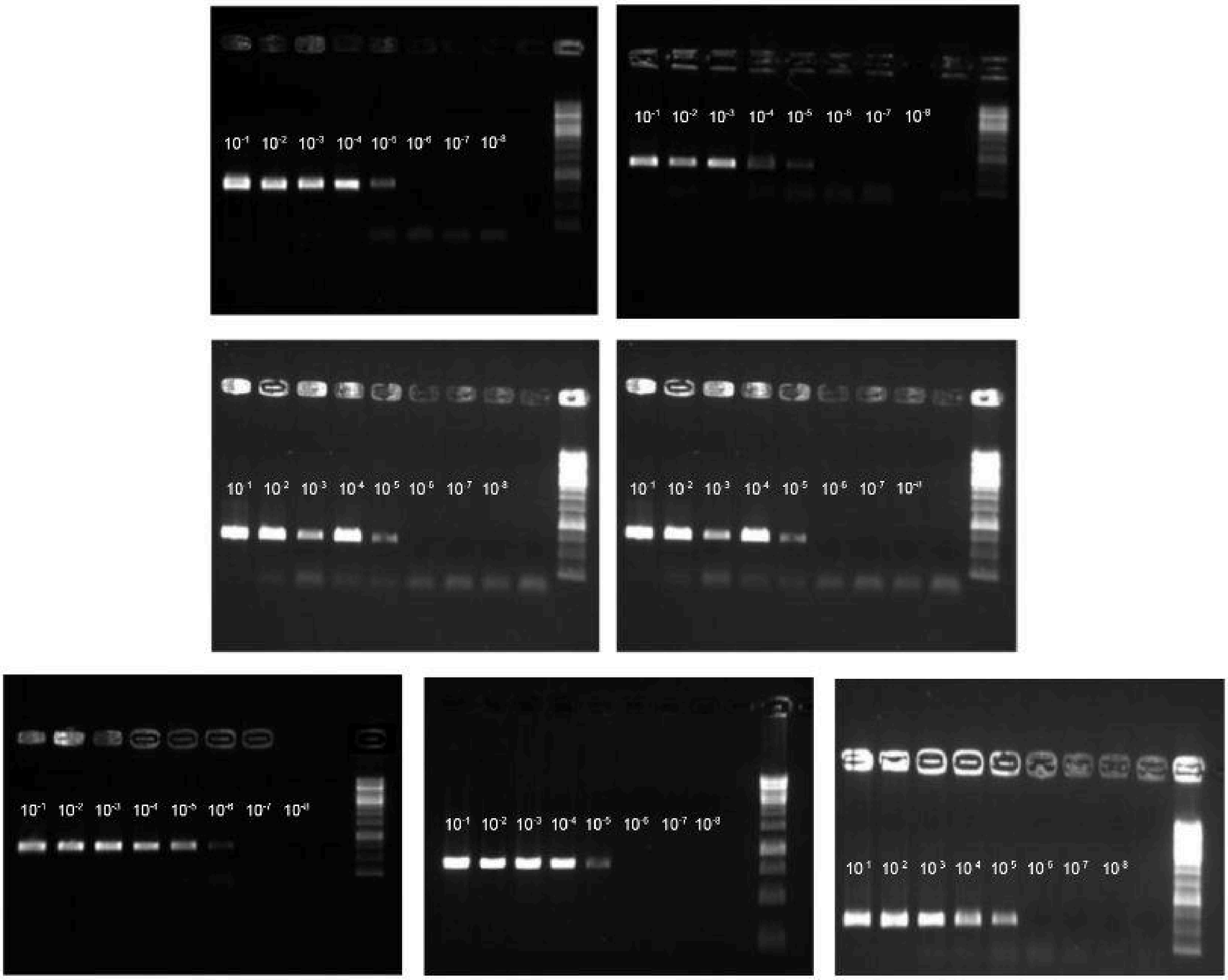

**Supplementary Figure 2.** Results of PCR performed for the set of primers designed for the new MLST scheme of *T. pallidum*. Each image corresponds to the following genes in the same order: *tp0136, tp0326, tp0548, tp0705, tp0858, tp0865, tp1031*. Likewise, for each gene, seven samples corresponding to a serial dilution of the Nichols sample were amplified, arranged in each image in the following order: Nichols, $10^{-1}$,$10^{-2}$,$10^{-3}$,$10^{-4}$,$10^{-5}$,$10^{-6}$,$10^{-7}$,$10^{-8}$ and the negative control.

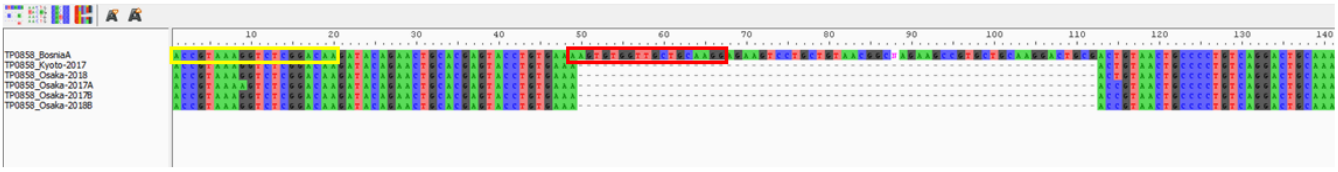

**Supplementary Figure 3.** The deletion found in the original *tp0858* primer for the five TEN samples from Japan. In yellow is highlighted the new forward primer designed for the *tp0858* gene and in pink the old one.

### REFERENCES

1. Giacani L, Iverson-Cabral SL, King JCK, Molini BJ, Lukehart SA, Centurion-Lara A. Complete Genome Sequence of the Treponema pallidum subsp. pallidum Sea81-4 Strain. Genome Announc. 2014;2: 333–314.

2. Vrbová E, Noda AA, Grillová L, Rodríguez I, Forsyth A, Oppelt J, et al. Whole genome sequences of Treponema pallidum subsp. endemicum isolated from Cuban patients: The non-clonal character of isolates suggests a persistent human infection rather than a single outbreak. PLoS Negl Trop Dis. 2022;16: e0009900.

3. Čejková D, Zobaníková M, Chen L, Pospíšilová P, Strouhal M, Qin X, et al. Whole Genome Sequences of Three Treponema pallidum ssp. pertenue Strains: Yaws and Syphilis Treponemes Differ in Less than 0.2% of the Genome Sequence. Lukehart S, editor. PLoS Negl Trop Dis. 2012;6: e1471.

4. Pětrošová H, Pospíšilová P, Strouhal M, Čejková D, Zobaníková M, Mikalová L, et al. Resequencing of Treponema pallidum ssp. pallidum strains Nichols and SS14: correction of sequencing errors resulted in increased separation of syphilis treponeme subclusters. PLoS One. 2013;8: e74319.

5. Arora N, Schuenemann VJ, Jäger G, Peltzer A, Seitz A, Herbig A, et al. Origin of modern syphilis and emergence of a pandemic Treponema pallidum cluster. Nat Microbiol. 2016;2: 16245.

6. Strouhal M, Mikalová L, Havlíčková P, Tenti P, Čejková D, Rychlík I, et al. Complete genome sequences of two strains of Treponema pallidum subsp. pertenue from Ghana, Africa: Identical genome sequences in samples isolated more than 7 years apart. PLoS Negl Trop Dis. 2017;11: e0005894.

7. Grillová L, Oppelt J, Mikalová L, Nováková M, Giacani L, Niesnerová A, et al. Directly Sequenced Genomes of Contemporary Strains of Syphilis Reveal Recombination-Driven Diversity in Genes Encoding Predicted Surface-Exposed Antigens. Front Microbiol. 2019;10: 1691.

8. Strouhal M, Mikalová L, Haviernik J, Knauf S, Bruisten S, Noordhoek GT, et al. Complete genome sequences of two strains of Treponema pallidum subsp. pertenue from Indonesia: Modular structure of several treponemal genes. PLoS Negl Trop Dis. 2018;12: e0006867.

9. Grillová L, Bawa T, Mikalová L, Gayet-Ageron A, Nieselt K, Strouhal M, et al. Molecular characterization of Treponema pallidum subsp. pallidum in Switzerland and France with a new multilocus sequence typing scheme. PLoS One. 2018;13: e0200773.

10. Katz SS, Chi K-H, Nachamkin E, Danavall D, Taleo F, Kool JL, et al. Molecular strain typing of the yaws pathogen, Treponema pallidum subspecies pertenue. Kalendar R, editor. PLoS One. 2018;13: e0203632.

11. Chuma IS, Roos C, Atickem A, Bohm T, Anthony Collins D, Grillová L, et al. Strain diversity of Treponema pallidum subsp. pertenue suggests rare interspecies transmission in African nonhuman primates. Sci Rep. 2019;9: 14243.

12. Medappa M, Pospíšilová P, Madruga MPM, John LN, Beiras CG, Grillová L, et al. Low genetic diversity of Treponema pallidum ssp. pertenue (TPE) isolated from patients' ulcers in Namatanai District of Papua New Guinea: Local human population is infected by three TPE

genotypes. PLoS Negl Trop Dis. 2024;18: e0011831.

13. Vaulet LG, Grillová L, Mikalová L, Casco R, Fermepin MR, Pando MA, et al. Molecular typing of treponema pallidum isolates from buenos aires, Argentina: Frequent nichols-like isolates and low levels of macrolide resistance. PLoS One. 2017. doi:10.1371/journal.pone.0172905

14. Mikalová L, Grillová L, Osbak K, Strouhal M, Kenyon C, Crucitti T, et al. Molecular Typing of Syphilis-Causing Strains Among Human Immunodeficiency Virus-Positive Patients in Antwerp, Belgium. Sex Transm Dis. 2017;44: 376–379.

15. Flores JA, Vargas SK, Leon SR, Perez DG, Ramos LB, Chow J, et al. Treponema pallidum pallidum Genotypes and Macrolide Resistance Status in Syphilitic Lesions among Patients at 2 Sexually Transmitted Infection Clinics in Lima, Peru. Sex Transm Dis. 2016;43: 465–466.

16. Wu H, Chang S-Y, Lee N-Y, Huang W-C, Wu B-R, Yang C-J, et al. Evaluation of macrolide resistance and enhanced molecular typing of Treponema pallidum in patients with syphilis in Taiwan: a prospective multicenter study. J Clin Microbiol. 2012;50: 2299–2304.

17. Read P, Tagg KA, Jeoffreys N, Guy RJ, Gilbert GL, Donovan B. Treponema pallidum Strain Types and Association with Macrolide Resistance in Sydney, Australia: New TP0548 Gene Types Identified. J Clin Microbiol. 2016;54: 2172–2174.

18. Noda AA, Matos N, Blanco O, Rodríguez I, Stamm LV. First Report of the 23S rRNA Gene A2058G Point Mutation Associated With Macrolide Resistance in Treponema pallidum From Syphilis Patients in Cuba. Sex Transm Dis. 2016;43: 332–334.

19. Xiao Y, Liu S, Liu Z, Xie Y, Jiang C, Xu M, et al. Molecular Subtyping and Surveillance of Resistance Genes In Treponema pallidum DNA From Patients With Secondary and Latent Syphilis in Hunan, China. Sex Transm Dis. 2016;43: 310–316.

20. Zondag HCA, Bruisten SM, Vrbová E, Šmajs D. No bejel among Surinamese, Antillean and Dutch syphilis diagnosed patients in Amsterdam between 2006-2018 evidenced by multi-locus sequence typing of Treponema pallidum isolates. PLoS One. 2020;15: e0230288.

21. Vrbová E, Grillová L, Mikalová L, Pospíšilová P, Strnadel R, Dastychová E, et al. MLST typing of Treponema pallidum subsp. pallidum in the Czech Republic during 2004-2017: Clinical isolates belonged to 25 allelic profiles and harbored 8 novel allelic variants. PLoS One. 2019;14: e0217611.

22. Fernández-Naval C, Arando M, Espasa M, Antón A, Fernández-Huerta M, Silgado A, et al. Multilocus sequence typing of Treponema pallidum subsp. pallidum in Barcelona. Future Microbiol. 2021;16: 967–976.

23. Sahi SK, Zahlan JM, Tantalo LC, Marra CM. A Comparison of Treponema pallidum Subspecies pallidum Molecular Typing Systems: Multilocus Sequence Typing vs. Enhanced Centers for Disease Control and Prevention Typing. Sex Transm Dis. 2021;48: 670–674.

24. Garcia LN, Morando N, Otero AV, Moroni S, Moscatelli GF, Gonzalez N, et al. Multilocus sequence typing of Treponema pallidum pallidum in children with acquired syphilis by nonsexual contact. Future Microbiol. 2022;17: 1295–1305.

25. Venter JME, Müller EE, Mahlangu MP, Kularatne RS. Treponema pallidum Macrolide Resistance and Molecular Epidemiology in Southern Africa, 2008 to 2018. J Clin Microbiol.

2021;59: e0238520.

26. Pillay A, Lee M-K, Slezak T, Katz SS, Sun Y, Chi K-H, et al. Increased Discrimination of Treponema pallidum Strains by Subtyping With a 4-Component System Incorporating a Mononucleotide Tandem Repeat in rpsA. Sex Transm Dis. 2019;46: e42–e45.

27. Pospíšilová P, Grange PA, Grillová L, Mikalová L, Martinet P, Janier M, et al. Multi-locus sequence typing of Treponema pallidum subsp. pallidum present in clinical samples from France: Infecting treponemes are genetically diverse and belong to 18 allelic profiles. Marangoni A, editor. PLoS One. 2018;13: e0201068.

28. Morando N, Vrbová E, Melgar A, Rabinovich RD, Šmajs D, Pando MA. High frequency of Nichols-like strains and increased levels of macrolide resistance in Treponema pallidum in clinical samples from Buenos Aires, Argentina. Sci Rep. 2022;12: 16339.

29. Sato W, Sedohara A, Koga M, Nakagama Y, Yotsuyanagi H, Kido Y, et al. Epidemic of multiple Treponema pallidum strains in men who have sex with men in Japan: efficient multi-locus sequence typing scheme and indicator biomarkers. AIDS Res Ther. 2024;21: 71.

30. Cummings OW, Durand ML, Barshak MB, Bispo PJM. Molecular Detection and Typing of in Non-Ocular Samples from Patients with Ocular Syphilis. Ocul Immunol Inflamm. 2024;32: 1580–1584.

31. Muhammad I, Khalifa EH, Salih MM, Ullah W, Elseid MSA, Qasim M, et al. Analysis of molecular subtypes and antibiotic resistance in Treponema pallidum isolates from blood donors in Khyber Pakhtunkhwa, Pakistan. PLoS One. 2024;19: e0305720.

32. Nadal-Barón P, Trejo-Zahínos J, Arando M, Barberan-Masegosa A, Bernat-Sole M, Pérez-Ugarte A, et al. High increase of Nichols-like clade circulating Treponema pallidum subsp. pallidum in Barcelona from 2021 to 2023. Sci Rep. 2024;14. doi:10.1038/s41598-024-74355-y

33. Vrbová E, Pospíšilová P, Dastychová E, Kojanová M, Kreidlová M, Rob F, et al. Majority of Treponema pallidum ssp. pallidum MLST allelic profiles in the Czech Republic (2004-2022) belong to two SS14-like clusters. Sci Rep. 2024;14: 17463.

34. Pillay A, Vilfort K, Debra A, Katz SS, Thurlow CM, Joseph SJ, et al. Molecular investigation of strains associated with ocular syphilis in the United States, 2016-2020. Microbiol Spectr. 2024;12: e0058124.

35. Esteves LS, Grassi VMT, Grassi LT, Nicola MRC, Silva MSN, Rossetti MLR, et al. Multilocus sequence typing of Treponema pallidum in male patients with genital ulcers in a public sexually transmitted infections clinic: a new allele and almost complete macrolide resistance. An Bras Dermatol. 2025. doi:10.1016/j.abd.2024.10.008

36. Imai K, Sato A, Tanaka M, Ohama Y, Nakayama S-I, Omachi R, et al. Prospective evaluation of non-invasive saliva specimens for the diagnosis of syphilis and molecular surveillance of. J Clin Microbiol. 2024;62: e0080924.

37. Queiroz JHF de S, Ferreira T da S, Lima BF, Perez EV de O, Mello CD de O, Simionatto S. Molecular characterization of Treponema pallidum isolates from Brazil. Diagn Microbiol Infect Dis. 2024;109: 116333.
