## Supplementary figures and images for "A new typing scheme demonstrates high discriminatory power for *Treponema pallidum* subspecies"

### Supplementary_figure1.png

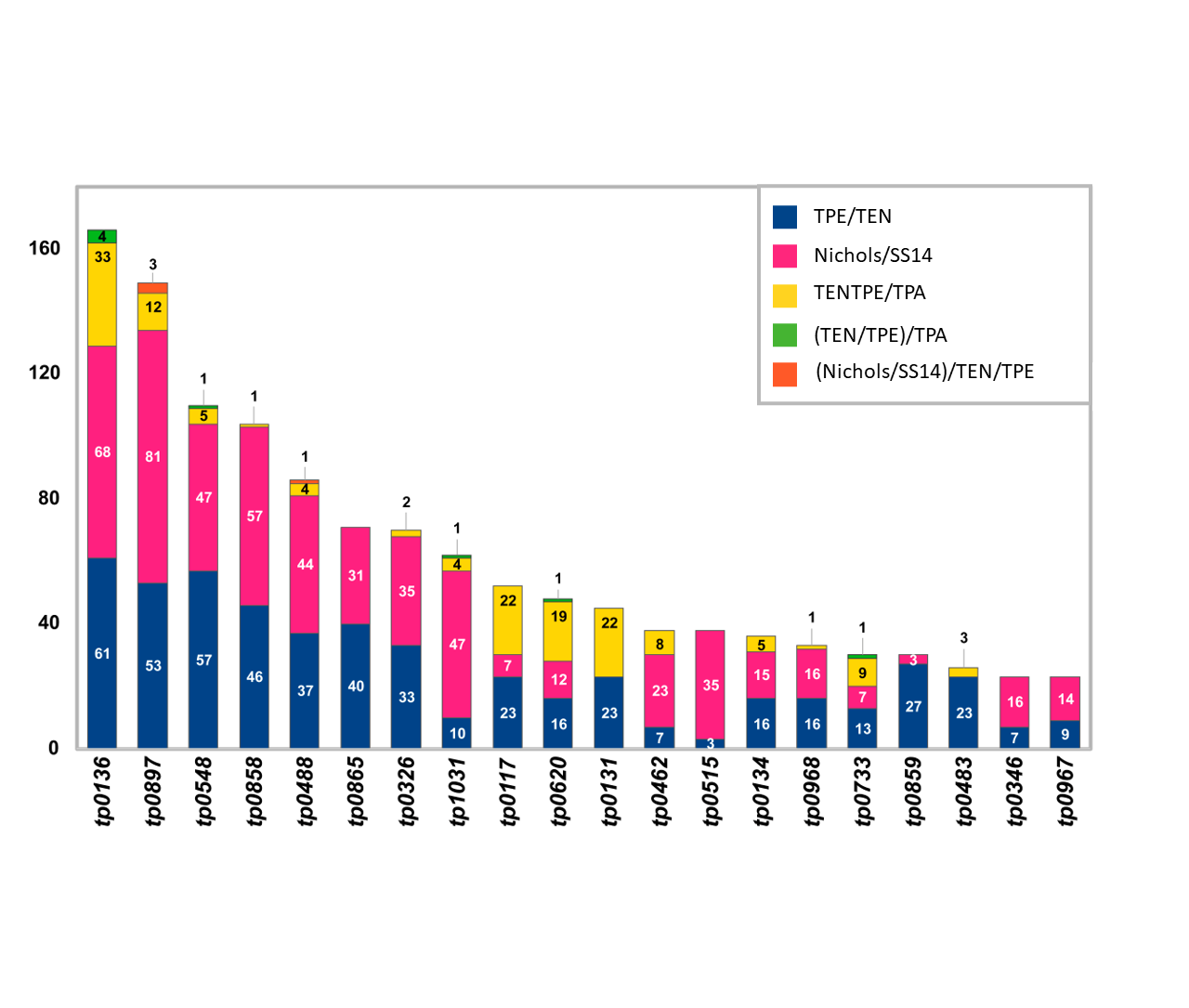

### Supplementary_figure3.png

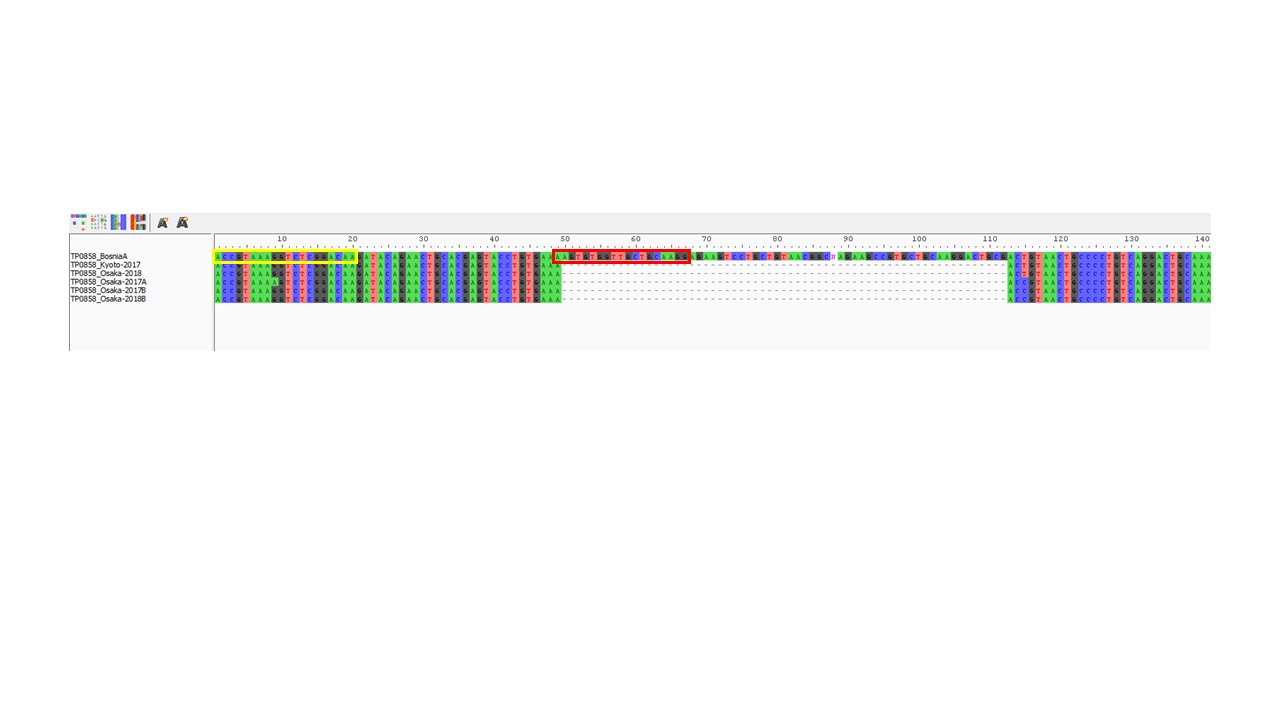

### Supplementary_figure3.pptx

#### Slide 1
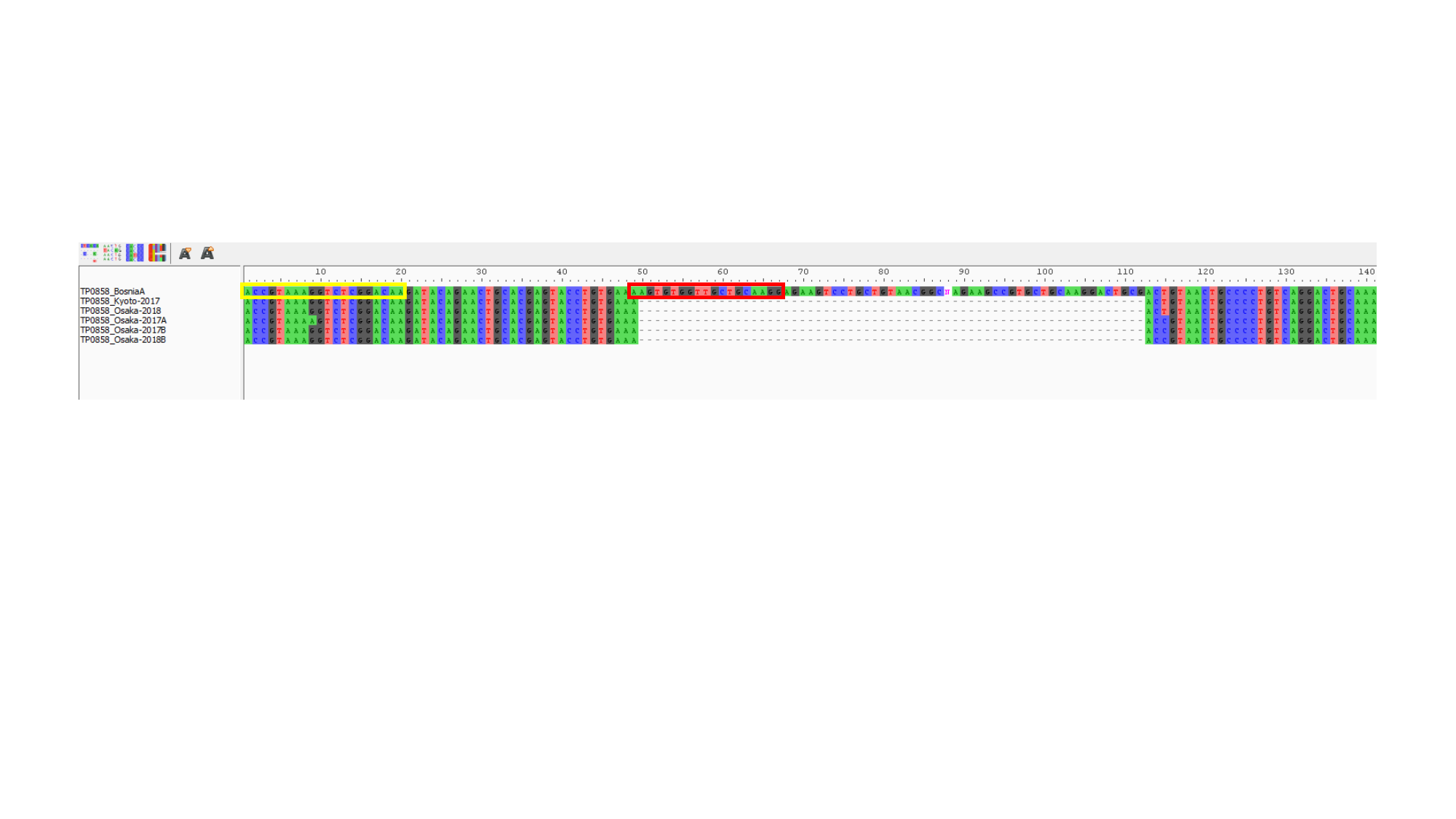
