## Supplementary_Files for "A new typing scheme demonstrates high discriminatory power for *Treponema pallidum* subspecies": Supplementary_File_1.py

```
#Marta Pla Diaz, Nov 7 (2019), for python2.7

#In-house script to compute the number of SNPs per gene
#Output: Fasta files per gene along with the folder route.

from os import listdir


def abrir_fichero(nombre_fichero):
    secuencias = []
    fich =  open(nombre_fichero, 'r')
    seq = ""
    for line in fich:
        if line.endswith('\n'):  
            line = line[:len(line) - 1]
        if line.endswith('\r'):  
            line = line[:len(line) - 1]
        if line.startswith(">"):
            if seq != "":
                secuencias.append(seq)
                seq = ""
                secuencias.append(line)
            else:
                secuencias.append(line)
        else:
            seq = seq + line
    secuencias.append(seq)
    return secuencias


def crear_diccionario(lista):
    diccionario = {}
    contador = 0
    while contador < len(lista):
        if lista[contador].startswith(">"):
            new_clave = str(lista[contador].replace(">", ""))
            diccionario[new_clave] = ""
            clave = new_clave
            contador = contador + 1
        else:
            diccionario[clave]= lista[contador]
            contador = contador + 1
    for genoma in diccionario:
        seq_genoma = str(diccionario[genoma])
        seq_genoma_upper = seq_genoma.upper()
        diccionario[genoma] = seq_genoma_upper
    return diccionario

def obtener_ref(dicc_seq):
    seq_ref = dicc_seq["NIC"]
    ref = "NIC"
    return seq_ref, ref

def anotar_cambios():
    dicc_cambios = {}
    return dicc_cambios

def procesar_seq(dicc_seq, dicc_cambios):
    for clave_other in dicc_seq:
        for clave_comp in dicc_seq:
            valor_other = str(dicc_seq[clave_other])
            valor_other_comp = str(dicc_seq[clave_comp])
            dicc_cambios = comparar(clave_other, valor_other, valor_other_comp, dicc_cambios)
    dicc_cambios_final = dicc_cambios
    num_cambios_total = len(dicc_cambios_final)
    return dicc_cambios_final, num_cambios_total

#changes the strain names accoding to your alignment 
def comprobar_subespecie(cepa):
    lista_yaws = "133_CDC2_map","AGU007_CDC2_map","Boe_92_CDC2_map","CDC2_real","CDC_2575_real","CHS119_CDC2_map",\
                 "SamoaD_real","NKNP1_CDC2_map","R8-KOS-007_CDC2_map","IND1_CDC2_map","133_CDC2_map",\
                 "WP0022.7.scab_CDC2_map","GHA1_CDC2_map","Fribourg_real","BAK_2117_CDC2_map","WP0022.7.liq_CDC2_map",\
                 "SJ003_CDC2_map","OKA_2116_CDC2_map","HATO_CDC2_map","Gauthier_real","K363_real","LMNP-1_real",\
                 "Boe_92_CDC2_map","CDC_2575_real","R3-SAL-007_CDC2_map","K403_real","CHS119_CDC2_map"
    lista_bejel = "BosniaA_real","Iraq B_real"
    lista_Nichols= "BAL3_Nichols_mapp","BAL73_Nichols_mapp","Sea_81-4_real","CW59__Nichols_mapp","CW65__Nichols_mapp",\
                   "CW82_Nichols_mapp","Nichols_real","CW86_Nichols_mapp","Chicago_real","PT_SIF1156_SS14_mapp",\
                   "CW83_Nichols_mapp","NL14_Nichols_mapp","BAL73_Nichols_mapp","W86_Nichols_mapp","NE20_Nichols_mapp",\
                   "SEA86_Nichols_mapp","CW82_Nichols_mapp","BAL3_Nichols_mapp","NIC2_Nichols_mapp"
    lista_SS14= "94A_SS14_mapp","94B_SS14_mapp","AR2_SS14_mapp","AU15_SS14_mapp","AU16_SS14_mapp","AU17_SS14_mapp",\
                "C3_SS14_mapp","CZ27_SS14_mapp","SW8_SS14_mapp","C3_SS14_mapp","UZ1974_SS14_mapp","NL11_SS14_mapp",\
                "UW824B_SS14_mapp","AU16_SS14_mapp","PT_SIF1242_SS14_mapp","UW370B_SS14_mapp","PT_SIF1020_SS14_mapp",\
                "NE15_SS14_mapp","MexicoA_real","AR2_SS14_mapp","W86_SS14_mapp","NL10_SS14_mapp","SS14_real",\
                "SHDR_SS14_mapp","CZ33_SS14_mapp","NE17_SS14_mapp","K3_SS14_mapp","CW85_SS14_mapp","94B_SS14_mapp",\
                "PT_SIF1196_SS14_mapp","PD28_SS14_mapp","CW84_SS14_mapp","AU17_SS14_mapp","NL16_SS14_mapp",\
                "SW4_SS14_mapp","SW6_SS14_mapp","NE19_SS14_mapp","SJ219_SS14_mapp","UW116B_SS14_mapp","P3_SS14_mapp",\
                "UW337B_SS14_mapp","GRA2_SS14_mapp","AU15_SS14_mapp","SHD-R_SS14_mapp",
    if cepa in lista_yaws:
        cepa_final = "Yaws"
    if cepa in lista_SS14:
        cepa_final = "SS14"
    if cepa in lista_Nichols:
        cepa_final = "Nichols"
    if cepa in lista_bejel:
        cepa_final = "Bejel"
    return cepa_final

def comparar(clave_other, valor_other, valor_other_comp,dicc_cambios):
    contador = 0
    while contador < len(valor_other_comp):
        if valor_other[contador] == valor_other_comp[contador]:
            contador += 1
        elif valor_other[contador] == "N" or valor_other[contador] == "-" or valor_other[contador] == "R" or \
                valor_other[contador] == "Y" or valor_other[contador] == "S" or valor_other[contador] == "M"\
                or valor_other_comp[contador] =="N"\
                or valor_other_comp[contador] =="-" or valor_other_comp[contador] =="R" or valor_other_comp[contador] == "Y"\
                or valor_other_comp[contador] == "S" or valor_other_comp[contador] == "M":
            contador += 1
        else:
            posicion = str(contador+1)
            valor_posi = str(valor_other[contador])
            if posicion in dicc_cambios:
                comprobacion = str(cepa_final + ":" + valor_posi)
                if comprobacion in dicc_cambios[posicion]:
                    contador += 1
                else:
                    dicc_cambios[posicion] = dicc_cambios[posicion] + ", "+ cepa_final + ":" + valor_posi
                    contador += 1
            else:
                dicc_cambios[posicion] =  cepa_final + ":" + valor_posi
                contador += 1
    return dicc_cambios


def ls(ruta = '.'):
    return listdir(ruta)

def main():
    lista_archivos = ls()
    for i in lista_archivos:
        if i.endswith(".fas"):
            nomfichero = str(i)
            secuencias = abrir_fichero(nomfichero)
            dicc_seq = crear_diccionario(secuencias)
            dicc_cambios = anotar_cambios()
            dicc_cambios_final, num_cambios_total = procesar_seq(dicc_seq, dicc_cambios)
            print "Numero total de SNPs en el gen " + nomfichero + ": " + str(num_cambios_total)
            
if __name__ == '__main__':
    main()
```
