## Supplementary_Files for "A new typing scheme demonstrates high discriminatory power for *Treponema pallidum* subspecies": Supplementary_File_2.py

```
##Marta Pla Diaz, Feb 28 (2019), for python2.7

# -*- coding: utf-8 -*-
#output: gene fasta files along with the folder route.

from os import listdir

def abrir_fichero(nombre_fichero):
    secuencias = []
    fich =  open(nombre_fichero, 'r')
    seq = ""
    for line in fich:
        if line.endswith('\n'):  
            line = line[:len(line) - 1]
        if line.endswith('\r'):  
            line = line[:len(line) - 1]
        if line.startswith(">"):
            if seq != "":
                secuencias.append(seq)
                seq = ""
                secuencias.append(line)
            else:
                secuencias.append(line)
        else:
            seq = seq + line
    secuencias.append(seq)
    return secuencias


def crear_diccionario(lista):
    diccionario = {}
    contador = 0
    while contador < len(lista):
        if lista[contador].startswith(">"):
            new_clave = str(lista[contador].replace(">", ""))
            diccionario[new_clave] = ""
            clave = new_clave
            contador = contador + 1
        else:
            diccionario[clave] = lista[contador]
            contador = contador + 1
    return diccionario

def obtener_ref(dicc_seq):
    seq_ref = dicc_seq["NIC"]
    ref = "NIC"
    return seq_ref, ref

def anotar_cambios():
    dicc_cambios = {}
    return dicc_cambios

def procesar_seq(dicc_seq, seq_ref, dicc_cambios):
    for clave_other in dicc_seq:
        valor_other = str(dicc_seq[clave_other])
        dicc_cambios = comparar(clave_other, valor_other, seq_ref, dicc_cambios)
    dicc_cambios_final = dicc_cambios
    num_cambios_total = len(dicc_cambios_final)
    return dicc_cambios_final, num_cambios_total

#change to your strain list
def comprobar_subespecie(cepa):
    lista_yaws = "SAM1","SAMD","CDC2","IND1" ,"FRI","GAU","GHA1","CDC2575","GHANA_051","SI30","SI28","SI17","SI03",\
                 "SI32","SI20","SI37","JK3168"
    lista_bejel = "BOSA","Cuba_60_I","Cuba_60_II"
    lista_Nichols= "NE20", "AU9","CHI","NIC","NIC1", "DAL", "NIC2","BAL3","SEA81","SEA86","UW189B",\
                   "BAL73","DZH7", "SEA81_4_2","SEA81_4_1"
    lista_SS14= "Cuba_178","UW228B","Q3","PT_SIF0697","CZ33","PT_SIF1142","PT_SIF1020","PT_SIF1140","PT_SIF1261",\
                "A2","GRA2","AR2","DZH3","SHD-R","PT_SIF1278","Amoy","PT_SIF1156","B3","Cuba_Vi2","NE17","NE19","NE12",\
                "NE13","NE14","NE15","AU17","Cuba_55_I","PT_SIF0875","AU15","AU16","PT_SIF0751","AU13","K3",\
                "PT_SIF1167","PT_SIF1200","SW1","SW2","C3","PT_SIF1280","SW4","SW6","UW391B","SW8","Cuba_167_I",\
                "PT_SIF0908","UW254B","SHC_O","PT_SIF1299","AU4","Cuba_68_II","Cali_167","PT_SIF1348","PT_SIF1063",\
                "PT_SIF1183","PT_SIF0877_3","A2_2","UW074B","SHG_I2","Cali_156","Cuba_120","Cuba_189_II","MEXA","SS14",\
                "PT_SIF1196","IMD6","IMD4","IMD5","Cuba_113","PT_SIF0954","PT_SIF1127","PT_SIF1002","IMD24",\
                "PT_SIF1242","UW1","IMD29","CZ27","P3","PT_SIF1135","SHE_V","PT_SIF1252","IMD17","Cuba_49_II","DZH11",\
                "DZH16", "JK3167","JK3170"
    if cepa in lista_yaws:
        cepa_final = "Yaws"
    if cepa in lista_SS14:
        cepa_final = "SS14"
    if cepa in lista_Nichols:
        cepa_final = "Nichols"
    if cepa in lista_bejel:
        cepa_final = "Bejel"
    return cepa_final

def comparar(clave_other, valor_other, seq_ref, dicc_cambios):
    contador = 0
    while contador < len(seq_ref):
        cepa_final = comprobar_subespecie(clave_other)
        if valor_other[contador] == seq_ref[contador]:
            contador += 1
        elif valor_other[contador] == "N" or valor_other[contador] == "-" or valor_other[contador] == "R" or \
                valor_other[contador] == "Y" or valor_other[contador] == "S" or valor_other[contador] == "M"\
            contador += 1
        else:
            posicion = str(contador)
            valor_posi = str(valor_other[contador])
            if posicion in dicc_cambios:
                comprobacion = str(cepa_final + ":" + valor_posi)
                if comprobacion in dicc_cambios[posicion]:
                    contador += 1
                else:
                    dicc_cambios[posicion] = dicc_cambios[posicion] + ", "+ cepa_final + ":" + valor_posi
                    contador += 1
            else:
                dicc_cambios[posicion] =  cepa_final + ":" + valor_posi
                comprobar = str("Otras" + ":" + seq_ref[contador])
                if not comprobar in dicc_cambios[posicion]:
                    dicc_cambios[posicion] = dicc_cambios[posicion] + ", " + "Otras" + ":" + seq_ref[contador]
                contador += 1
    return dicc_cambios

def diccionario_final_bien(dicc_cambios_final,dicc_cambios_final2, numseq):
    diccionario_subespecies_cambi = {}
    dicc_cambios_final_ordenado = sorted(dicc_cambios_final.items(),key=lambda x: int(x[0]),)
    for i in dicc_cambios_final_ordenado:
        posicion = i[0]
        if posicion in dicc_cambios_final2:
            cepas_contar = dicc_cambios_final2[posicion]
            yaws_final, bejel_final, SS14_final, Nichols_final, num_otras = obtener_contadores_ycepasfinal(cepas_contar, numseq)
        subespecie = str(i[1])
        lista_subespecie = subespecie.split(",")
        sub_bejel = ""
        sub_Nichols = ""
        sub_yaws = ""
        sub_SS14 = ""
        sub_otras = ""
        for j in lista_subespecie:
            if "Bejel" in j:
                sub_bejel= bejel_final
            if "Yaws" in j:
                sub_yaws = yaws_final
            if "Nichols" in j:
                sub_Nichols = Nichols_final
            if "SS14" in j:
                sub_SS14= SS14_final
            if "Otras" in j:
                sub_otras = j
        posicion_final = int(i[0]) + 1
        valor_dicc = sub_bejel + sub_Nichols + sub_yaws +  sub_SS14 + sub_otras
        diccionario_subespecies_cambi[posicion_final] = valor_dicc
        diccionario_subespecies_cambi
    return diccionario_subespecies_cambi

def obtener_contadores_ycepasfinal(cepas_contar, numseq):
    lista_cepas = cepas_contar.split(",")
    contador_yaws = 0
    contador_SS14 = 0
    contador_Nichols = 0
    contador_bejel = 0
    lista_cambios_yaws = ""
    lista_cambios_bejel = ""
    lista_cambios_SS14 = ""
    lista_cambios_Nichols = ""
    yaws_final = ""
    bejel_final = ""
    SS14_final = ""
    Nichols_final = ""
    for cepa in lista_cepas:
        obtecambio_cepa = cepa.split(":")
        cambio_cepa = str(obtecambio_cepa[1])
        cepa_sincambio = cepa[:len(cepa) - 2]
        cepa_final_def= comprobar_subespecie2(cepa_sincambio)
        if cepa_final_def == "Yaws":
            lista_cambios_yaws = lista_cambios_yaws + cambio_cepa
        if cepa_final_def == "Bejel":
            lista_cambios_bejel = lista_cambios_bejel + cambio_cepa
        if cepa_final_def == "SS14":
            lista_cambios_SS14 = lista_cambios_SS14 + cambio_cepa
        if cepa_final_def == "Nichols":
            lista_cambios_Nichols = lista_cambios_Nichols + cambio_cepa
    lista_cambios_yaws_set = set(lista_cambios_yaws)
    lista_cambios_bejel_set = set(lista_cambios_bejel)
    lista_cambios_SS14_set = set(lista_cambios_SS14)
    lista_cambios_Nichols_set = set(lista_cambios_Nichols)

    if len(lista_cambios_yaws_set) == 1:
        contador_yaws = len(lista_cambios_yaws)
        yaws_final = "Yaws" + ":" + lista_cambios_yaws[0] + ","
    if len(lista_cambios_yaws_set) == 2:
        element1 = tuple(lista_cambios_yaws_set)[0]
        element2 = tuple(lista_cambios_yaws_set)[1]
        contadoryaws1 =  "Yaws" + ":" + element1 + ","
        contadoryaws2 =  "Yaws" + ":" + element2 + ","
        yaws_final = contadoryaws1 + contadoryaws2
        contador_yaws = lista_cambios_yaws.count(element1) + lista_cambios_yaws.count(element2)
    if len(lista_cambios_yaws_set) == 3:
        element1 = tuple(lista_cambios_yaws_set)[0]
        element2 = tuple(lista_cambios_yaws_set)[1]
        element3 = tuple(lista_cambios_yaws_set)[2]
        contadoryaws1 = "Yaws" + ":" + element1 + ","
        contadoryaws2 = "Yaws" + ":" + element2 + ","
        contadoryaws3 = "Yaws" + ":" + element3 + ","
        yaws_final = contadoryaws1 + contadoryaws2 + contadoryaws3
        contador_yaws = lista_cambios_yaws.count(element1) + lista_cambios_yaws.count(element2) + lista_cambios_yaws.count(element3)
    if len(lista_cambios_yaws_set) == 3:
        element1 = tuple(lista_cambios_yaws_set)[0]
        element2 = tuple(lista_cambios_yaws_set)[1]
        element3 = tuple(lista_cambios_yaws_set)[2]
        element4 = tuple(lista_cambios_yaws_set)[3]
        contadoryaws1 = "Yaws" + ":" + element1 + ","
        contadoryaws2 = "Yaws" + ":" + element2 + ","
        contadoryaws3 = "Yaws" + ":" + element3 + ","
        contadoryaws4 = "Yaws" + ":" + element4 + ","
        yaws_final = contadoryaws1 + contadoryaws2 + contadoryaws3 + contadoryaws4
        contador_yaws = lista_cambios_yaws.count(element1) + lista_cambios_yaws.count(element2) +\
                        lista_cambios_yaws.count(element3) + lista_cambios_yaws.count(element4)

    if len(lista_cambios_bejel_set) == 1:
        contador_bejel = len(lista_cambios_bejel)
        bejel_final = "Bejel" + ":" + lista_cambios_bejel[0] + ","
    if len(lista_cambios_bejel_set) == 2:
        element1 = tuple(lista_cambios_bejel_set)[0]
        element2 = tuple(lista_cambios_bejel_set)[1]
        contadorbejel1 = "Bejel" + ":" + element1 + ","
        contadorbejel2 = "Bejel" + ":" + element2 + ","
        bejel_final = contadorbejel1 + contadorbejel2
        contador_bejel = lista_cambios_bejel.count(element1) + lista_cambios_bejel.count(element2)
    if len(lista_cambios_bejel_set) == 3:
        element1 = tuple(lista_cambios_bejel_set)[0]
        element2 = tuple(lista_cambios_bejel_set)[1]
        element3 = tuple(lista_cambios_bejel_set)[2]
        contadorbejel1 = "Bejel" + ":" + element1 + ","
        contadorbejel2 = "Bejel" + ":" + element2 + ","
        contadorbejel3 = "Bejel" + ":" + element3 + ","
        bejel_final = contadorbejel1 + contadorbejel2 + contadorbejel3
        contador_bejel = lista_cambios_bejel.count(element1) + lista_cambios_bejel.count(element2) + lista_cambios_bejel.count(element3)
    if len(lista_cambios_bejel_set) == 4:
        element1 = tuple(lista_cambios_bejel_set)[0]
        element2 = tuple(lista_cambios_bejel_set)[1]
        element3 = tuple(lista_cambios_bejel_set)[2]
        element4 = tuple(lista_cambios_bejel_set)[3]
        contadorbejel1 = "Bejel" + ":" + element1 + ","
        contadorbejel2 = "Bejel" + ":" + element2 + ","
        contadorbejel3 = "Bejel" + ":" + element3 + ","
        contadorbejel4 = "Bejel" + ":" + element4 + ","
        bejel_final = contadorbejel1 + contadorbejel2 + contadorbejel3 + contadorbejel4
        contador_bejel = lista_cambios_bejel.count(element1) + lista_cambios_bejel.count(element2) +\
                         lista_cambios_bejel.count(element3) + lista_cambios_bejel.count(element4)

    if len(lista_cambios_SS14_set) == 1:
        contador_SS14 = len(lista_cambios_SS14)
        SS14_final = "SS14" + ":" + lista_cambios_SS14[0] + ","
    if len(lista_cambios_SS14_set) == 2:
        element1 = tuple(lista_cambios_SS14_set)[0]
        element2 = tuple(lista_cambios_SS14_set)[1]
        contadorSS141 = "SS14" + ":" + element1 + ","
        contadorSS142 = "SS14" + ":" + element2 + ","
        SS14_final = contadorSS141 + contadorSS142
        contador_SS14 = lista_cambios_SS14.count(element1) + lista_cambios_SS14.count(element2)
    if len(lista_cambios_SS14_set) == 3:
        element1 = tuple(lista_cambios_SS14_set)[0]
        element2 = tuple(lista_cambios_SS14_set)[1]
        element3 = tuple(lista_cambios_SS14_set)[2]
        contadorSS141 = "SS14" + ":" + element1 + ","
        contadorSS142 = "SS14" + ":" + element2 + ","
        contadorSS143 = "SS14" + ":" + element3 + ","
        SS14_final = contadorSS141 + contadorSS142 + contadorSS143
        contador_SS14 = lista_cambios_SS14.count(element1) + lista_cambios_SS14.count(element2) + lista_cambios_SS14.count(element3)
    if len(lista_cambios_SS14_set) == 4:
        element1 = tuple(lista_cambios_SS14_set)[0]
        element2 = tuple(lista_cambios_SS14_set)[1]
        element3 = tuple(lista_cambios_SS14_set)[2]
        element4 = tuple(lista_cambios_SS14_set)[3]
        contadorSS141 = "SS14" + ":" + element1 + ","
        contadorSS142 = "SS14" + ":" + element2 + ","
        contadorSS143 = "SS14" + ":" + element3 + ","
        contadorSS144 = "SS14" + ":" + element4 + ","
        SS14_final = contadorSS141 + contadorSS142 + contadorSS143 + contadorSS144
        contador_SS14 = lista_cambios_SS14.count(element1) + lista_cambios_SS14.count(element2) +\
                        lista_cambios_SS14.count(element3) + lista_cambios_SS14.count(element4)

    if len(lista_cambios_Nichols_set) == 1:
        contador_Nichols = len(lista_cambios_Nichols)
        Nichols_final = "Nichols" + ":" + lista_cambios_Nichols[0] + ","
    if len(lista_cambios_Nichols_set) == 2:
        element1 = tuple(lista_cambios_Nichols_set)[0]
        element2 = tuple(lista_cambios_Nichols_set)[1]
        contadorNichols1 = "Nichols" + ":" + element1 + ","
        contadorNichols2 = "Nichols" + ":" + element2 + ","
        Nichols_final = contadorNichols1  + contadorNichols2
        contador_Nichols = lista_cambios_Nichols.count(element1) + lista_cambios_Nichols.count(element2)
    if len(lista_cambios_Nichols_set) == 3:
        element1 = tuple(lista_cambios_Nichols_set)[0]
        element2 = tuple(lista_cambios_Nichols_set)[1]
        element3 = tuple(lista_cambios_Nichols_set)[2]
        contadorNichols1 = "Nichols" + ":" + element1 + ","
        contadorNichols2 = "Nichols" + ":" + element2 + ","
        contadorNichols3 = "Nichols" + ":" + element3 + ","
        Nichols_final = contadorNichols1 + contadorNichols2 + contadorNichols3
        contador_Nichols = lista_cambios_Nichols.count(element1) + lista_cambios_Nichols.count(element2) + lista_cambios_Nichols.count(element3)
    if len(lista_cambios_Nichols_set) == 3:
        element1 = tuple(lista_cambios_Nichols_set)[0]
        element2 = tuple(lista_cambios_Nichols_set)[1]
        element3 = tuple(lista_cambios_Nichols_set)[2]
        element4 = tuple(lista_cambios_Nichols_set)[3]
        contadorNichols1 = "Nichols" + ":" + element1 + ","
        contadorNichols2 = "Nichols" + ":" + element2 + ","
        contadorNichols3 = "Nichols" + ":" + element3 + ","
        contadorNichols4 = "Nichols" + ":" + element4 + ","
        Nichols_final = contadorNichols1 + contadorNichols2 + contadorNichols3 + contadorNichols4
        contador_Nichols = lista_cambios_Nichols.count(element1) + lista_cambios_Nichols.count(element2)\
                           + lista_cambios_Nichols.count(element3) + lista_cambios_Nichols.count(element4)

    suma_contadores =  contador_yaws + contador_bejel + contador_SS14 + contador_Nichols
    num_otras = numseq - suma_contadores
    return yaws_final, bejel_final, SS14_final, Nichols_final, num_otras


def anotar_cambios2():
    dicc_cambios2 = {}
    return dicc_cambios2

def procesar_seq_2(dicc_seq, seq_ref, dicc_cambios2):
    for clave_other in dicc_seq:
        valor_other = str(dicc_seq[clave_other])
        dicc_cambios2 = comparar2(clave_other, valor_other, seq_ref, dicc_cambios2)
        dicc_cambios_final2 = dicc_cambios2
    return dicc_cambios_final2

def comparar2(clave_other, valor_other, seq_ref, dicc_cambios2):
    contador = 0
    while contador < len(seq_ref):
        if valor_other[contador] == seq_ref[contador]:
            contador += 1
        elif valor_other[contador] == "N":
            contador += 1
        else:
            posicion = str(contador)
            valor_posi = str(valor_other[contador])
            if posicion in dicc_cambios2:
                dicc_cambios2[posicion] = dicc_cambios2[posicion] + ","+ clave_other + ":" + valor_posi
                contador += 1
            else:
                dicc_cambios2[posicion] =  clave_other + ":" + valor_posi
                contador += 1
    return dicc_cambios2

#change to your strain list
def comprobar_subespecie2(cepa):
    lista_yaws = "SAM1","SAMD","CDC2","IND1" ,"FRI","GAU","GHA1","CDC2575","GHANA_051","SI30","SI28","SI17","SI03",\
                 "SI32","SI20","SI37","JK3168"
    lista_bejel = "BOSA","Cuba_60_I","Cuba_60_II"
    lista_Nichols= "NE20", "AU9","CHI","NIC","NIC1", "DAL", "NIC2","BAL3","SEA81","SEA86","UW189B",\
                   "BAL73","DZH7", "SEA81_4_2","SEA81_4_1"
    lista_SS14= "Cuba_178","UW228B","Q3","PT_SIF0697","CZ33","PT_SIF1142","PT_SIF1020","PT_SIF1140","PT_SIF1261",\
                "A2","GRA2","AR2","DZH3","SHD-R","PT_SIF1278","Amoy","PT_SIF1156","B3","Cuba_Vi2","NE17","NE19","NE12",\
                "NE13","NE14","NE15","AU17","Cuba_55_I","PT_SIF0875","AU15","AU16","PT_SIF0751","AU13","K3",\
                "PT_SIF1167","PT_SIF1200","SW1","SW2","C3","PT_SIF1280","SW4","SW6","UW391B","SW8","Cuba_167_I",\
                "PT_SIF0908","UW254B","SHC_O","PT_SIF1299","AU4","Cuba_68_II","Cali_167","PT_SIF1348","PT_SIF1063",\
                "PT_SIF1183","PT_SIF0877_3","A2_2","UW074B","SHG_I2","Cali_156","Cuba_120","Cuba_189_II","MEXA","SS14",\
                "PT_SIF1196","IMD6","IMD4","IMD5","Cuba_113","PT_SIF0954","PT_SIF1127","PT_SIF1002","IMD24",\
                "PT_SIF1242","UW1","IMD29","CZ27","P3","PT_SIF1135","SHE_V","PT_SIF1252","IMD17","Cuba_49_II","DZH11",\
                "DZH16", "JK3167","JK3170"
    if str(cepa) in lista_yaws:
        cepa_final = "Yaws"
    if str(cepa) in lista_SS14:
        cepa_final = "SS14"
    if str(cepa) in lista_Nichols:
        cepa_final = "Nichols"
    if str(cepa) in lista_bejel:
        cepa_final = "Bejel"
    return cepa_final


def contar_cambios_interintra(diccionario_subespecies_cambi, nomfichero):
    contadorYvsB = 0
    contadorNichols_SS14 = 0
    contador_intersubsp_YBvsSIF = 0
    contador_intersubsp_YvsBvsSIF = 0
    contador_intersubsp_TPAvsTENTPE = 0
    contador_NicvsSS14vsYB = 0
    contador_NicvsSS14vsYvsB = 0
    contador_yaws = 0
    contador_bejel = 0
    contador_Nichols = 0
    contador_SS14 = 0
    lista_yaws = ["M"]
    lista_bejel = ["M"]
    lista_Nichols = ["M"]
    lista_SS14 = ["M"]
    nomfichero = nomfichero + "_out_scritp_ramas"
    fich = open(nomfichero, 'w')
    for key_posicion in diccionario_subespecies_cambi:
        valor_posicion = diccionario_subespecies_cambi[key_posicion]
        subspycamb_pos = valor_posicion.split(",")
        print key_posicion
        for subespecie in subspycamb_pos:
            contadorYvsB = 0
            contadorNichols_SS14 = 0
            contador_intersubsp_YBvsSIF = 0
            contador_intersubsp_YvsBvsSIF = 0
            contador_intersubsp_TPAvsTENTPE = 0
            contador_NicvsSS14vsYB = 0
            if "Yaws" in subespecie:
                if lista_yaws[0] == "M":
                    lista_yaws = []
                    lista_yaws.append(subespecie[5])
                else:
                    if subespecie[5] in lista_yaws:
                        pass
                    else:
                        lista_yaws.append(subespecie[5])
                contador_yaws += 1
            if "Bejel" in subespecie:
                if lista_bejel[0] == "M":
                    lista_bejel = []
                    lista_bejel.append(subespecie[6])
                else:
                    if subespecie[6] in lista_bejel:
                        pass
                    else:
                        lista_bejel.append(subespecie[6])
                contador_bejel += 1
            if "Nichols" in subespecie:
                if lista_Nichols[0] == "M":
                    lista_Nichols = []
                    lista_Nichols.append(subespecie[8])
                else:
                    if subespecie[8] in lista_Nichols:
                        pass
                    else:
                        lista_Nichols.append(subespecie[8])
                contador_Nichols += 1
            if "SS14" in subespecie:
                if lista_SS14[0] == "M":
                    lista_SS14= []
                    lista_SS14.append(subespecie[5])
                else:
                    if subespecie[5] in lista_SS14:
                        pass
                    else:
                        lista_SS14.append(subespecie[5])
                contador_SS14 += 1


        if len(lista_yaws) == len(lista_bejel) and len(lista_yaws) == len(lista_Nichols) and len(lista_yaws) == len(lista_SS14):
            contyaws = 0
            contbejel = 0
            contnichols = 0
            contSS14 = 0
            contadorYvsB, contadorNichols_SS14, contador_intersubsp_YBvsSIF, contador_intersubsp_YvsBvsSIF, contador_intersubsp_TPAvsTENTPE,\
            contador_NicvsSS14vsYB, cont_NicvsSS14vsYvsB_IND = comprobacion_cambios_subespecies(lista_yaws, lista_bejel, lista_Nichols, lista_SS14, contyaws, contbejel,contnichols,contSS14,
                                      contadorYvsB, contadorNichols_SS14, contador_intersubsp_YBvsSIF, contador_intersubsp_YvsBvsSIF,
                                      contador_intersubsp_TPAvsTENTPE, contador_NicvsSS14vsYB, contador_NicvsSS14vsYvsB)

        else:
            print "INTENTANDO ARREGLAR ESTA MIERDA ---------------------------------------------------------------"
            if len(lista_Nichols) >= 1 and len(lista_yaws) >= 1 and len(lista_bejel) >= 1 and len(lista_SS14) >= 1:
                contador_veces = 0
                contYvsB_IND = 0
                contNichols_SS14_IND = 0
                cont_intersubsp_YBvsSIF_IND = 0
                cont__intersubsp_YvsBvsSIF_IND = 0
                cont_intersubsp_TPAvsTENTPE_IND = 0
                cont_NicvsSS14vsYB_IND = 0
                cont_NicvsSS14vsYvsB_IND = 0
                while contador_veces < len(lista_yaws):
                    contyaws = contador_veces
                    contbejel = 0
                    contnichols = 0
                    contSS14 = 0
                    contYvsB_IND, contNichols_SS14_IND, cont_intersubsp_YBvsSIF_IND, cont__intersubsp_YvsBvsSIF_IND, cont_intersubsp_TPAvsTENTPE_IND, \
                    cont_NicvsSS14vsYB_IND, cont_NicvsSS14vsYvsB_IND = comprobacion_cambios_subespecies(lista_yaws, lista_bejel, lista_Nichols,
                                                                              lista_SS14, contyaws, contbejel,
                                                                              contnichols, contSS14, contYvsB_IND,
                                                                              contNichols_SS14_IND,
                                                                              cont_intersubsp_YBvsSIF_IND,
                                                                              cont__intersubsp_YvsBvsSIF_IND,
                                                                              cont_intersubsp_TPAvsTENTPE_IND,
                                                                              cont_NicvsSS14vsYB_IND, cont_NicvsSS14vsYvsB_IND)
                    contador_veces += 1
                while contador_veces < len(lista_bejel):
                    contyaws = 0
                    contbejel = contador_veces
                    contnichols = 0
                    contSS14 = 0
                    contYvsB_IND, contNichols_SS14_IND, cont_intersubsp_YBvsSIF_IND, cont__intersubsp_YvsBvsSIF_IND, cont_intersubsp_TPAvsTENTPE_IND, \
                    cont_NicvsSS14vsYB_IND, cont_NicvsSS14vsYvsB_IND = comprobacion_cambios_subespecies(lista_yaws, lista_bejel, lista_Nichols,
                                                                              lista_SS14, contyaws, contbejel,
                                                                              contnichols, contSS14, contYvsB_IND,
                                                                              contNichols_SS14_IND,
                                                                              cont_intersubsp_YBvsSIF_IND,
                                                                              cont__intersubsp_YvsBvsSIF_IND,
                                                                              cont_intersubsp_TPAvsTENTPE_IND,
                                                                              cont_NicvsSS14vsYB_IND, cont_NicvsSS14vsYvsB_IND)
                    contador_veces += 1
                while contador_veces < len(lista_SS14):
                    contyaws = 0
                    contbejel = 0
                    contnichols = 0
                    contSS14 = contador_veces
                    contYvsB_IND, contNichols_SS14_IND, cont_intersubsp_YBvsSIF_IND, cont__intersubsp_YvsBvsSIF_IND, cont_intersubsp_TPAvsTENTPE_IND, \
                    cont_NicvsSS14vsYB_IND, cont_NicvsSS14vsYvsB_IND = comprobacion_cambios_subespecies(lista_yaws, lista_bejel, lista_Nichols,
                                                                              lista_SS14, contyaws, contbejel,
                                                                              contnichols, contSS14, contYvsB_IND,
                                                                              contNichols_SS14_IND,
                                                                              cont_intersubsp_YBvsSIF_IND,
                                                                              cont__intersubsp_YvsBvsSIF_IND,
                                                                              cont_intersubsp_TPAvsTENTPE_IND,
                                                                              cont_NicvsSS14vsYB_IND, cont_NicvsSS14vsYvsB_IND)
                    contador_veces += 1
                while contador_veces < len(lista_Nichols):
                    contyaws = 0
                    contbejel = 0
                    contnichols = contador_veces
                    contSS14 = 0
                    contYvsB_IND, contNichols_SS14_IND, cont_intersubsp_YBvsSIF_IND, cont__intersubsp_YvsBvsSIF_IND, cont_intersubsp_TPAvsTENTPE_IND, \
                    cont_NicvsSS14vsYB_IND, cont_NicvsSS14vsYvsB_IND = comprobacion_cambios_subespecies(lista_yaws, lista_bejel, lista_Nichols,
                                                                              lista_SS14, contyaws, contbejel,
                                                                              contnichols, contSS14, contYvsB_IND,
                                                                              contNichols_SS14_IND,
                                                                              cont_intersubsp_YBvsSIF_IND,
                                                                              cont__intersubsp_YvsBvsSIF_IND,
                                                                              cont_intersubsp_TPAvsTENTPE_IND,
                                                                              cont_NicvsSS14vsYB_IND, cont_NicvsSS14vsYvsB_IND)
                    contador_veces += 1
                if contYvsB_IND > 0:
                    contadorYvsB += 1
                if contNichols_SS14_IND > 0:
                    contadorNichols_SS14 += 1
                if cont_intersubsp_YBvsSIF_IND > 0:
                    contador_intersubsp_YBvsSIF += 1
                if cont__intersubsp_YvsBvsSIF_IND > 0:
                    contador_intersubsp_YvsBvsSIF += 1
                if cont_intersubsp_TPAvsTENTPE_IND > 0:
                    contador_intersubsp_TPAvsTENTPE += 1
                if cont_NicvsSS14vsYB_IND > 0:
                    contador_NicvsSS14vsYB += 1
                if cont_NicvsSS14vsYvsB_IND >0:
                    contador_NicvsSS14vsYvsB += 1

        suma = contador_intersubsp_TPAvsTENTPE + contador_intersubsp_YBvsSIF
        print suma, contador_intersubsp_TPAvsTENTPE, contador_intersubsp_YBvsSIF

        fich.write(str(key_posicion) + " " + str(contadorYvsB) + " " + str(contadorNichols_SS14) + " " + str(suma) + " "
                   +str(contador_intersubsp_YvsBvsSIF)+ " " + str(contador_NicvsSS14vsYB) + " " + str(contador_NicvsSS14vsYvsB) + "\n")

        print valor_posicion
        print contadorYvsB, "contadorYvsB"
        print contadorNichols_SS14,  "contadorNichols_SS14"
        print contador_intersubsp_YBvsSIF, "contador_intersubsp_YBvsSIF"
        print contador_intersubsp_YvsBvsSIF, "contador_intersubsp_YvsBvsSI"
        print contador_intersubsp_TPAvsTENTPE, "contador_intersubsp_TPAvsTENTPE"
        print contador_NicvsSS14vsYB, "contador_NicvsSS14vsYB"
        print contador_NicvsSS14vsYvsB, "contador NicvsSS14vsYvsB "
        print "________________________________________________"

        lista_yaws = ["M"]
        lista_bejel = ["M"]
        lista_Nichols = ["M"]
        lista_SS14 = ["M"]


    return


def comprobacion_cambios_subespecies (lista_yaws, lista_bejel, lista_Nichols, lista_SS14, contyaws, contbejel,contnichols,contSS14,
                                      contadorYvsB, contadorNichols_SS14, contador_intersubsp_YBvsSIF, contador_intersubsp_YvsBvsSIF,
                                      contador_intersubsp_TPAvsTENTPE, contador_NicvsSS14vsYB, contador_NicvsSS14vsYvsB):

    if lista_yaws[contyaws] != "M" and lista_yaws[contyaws] != lista_bejel[contbejel] or lista_bejel[contbejel] != "M" and lista_bejel[contbejel] != \
            lista_yaws[contyaws]:
        contadorYvsB += 1
        print "cambio TEN_TPE"
   
    if lista_Nichols[contnichols] != "M" and lista_Nichols[contnichols] != lista_SS14[contSS14] or lista_SS14[contSS14] != "M" and lista_SS14[contSS14] != \
            lista_Nichols[contnichols]:
        contadorNichols_SS14 += 1
        print "cambio NIC_SS14"
   
    if lista_yaws[contyaws] != "M" and lista_yaws[contyaws] == lista_bejel[contbejel] and lista_yaws[contyaws] != lista_Nichols[contnichols] and lista_yaws[
        contyaws] != lista_SS14[contSS14]:
        contador_intersubsp_YBvsSIF += 1
        print "mismo cambio TEN/TPE y diferentes de TPA"
    if lista_yaws[contyaws] != "M" and lista_bejel[contbejel] != "M" and lista_yaws[contyaws] != lista_bejel[contbejel] and lista_yaws[contyaws] != \
            lista_Nichols[contnichols] and lista_yaws[contyaws] != lista_SS14[contSS14] \
            and lista_bejel[contbejel] != lista_Nichols[contnichols] and lista_bejel[contbejel] != lista_SS14[contSS14]:
        contador_intersubsp_YvsBvsSIF += 1
        print "cambio difernente en TEN/TPE y ademas diferentes de TPA"
    
    if lista_Nichols[contnichols] != "M" and lista_Nichols[contnichols] == lista_SS14[contSS14] and lista_Nichols[contnichols] != lista_yaws[contyaws] and \
            lista_Nichols[0] != lista_bejel[contbejel]:
        contador_intersubsp_TPAvsTENTPE += 1
        print "Mismo cambio TPA y no en TEN ni TPE"
    if lista_Nichols[contnichols] != "M" and lista_SS14[contSS14] != "M" and lista_Nichols[contnichols] != lista_SS14[contSS14] and lista_Nichols[contnichols] != \
            lista_yaws[contyaws] and lista_Nichols[contnichols] != lista_bejel[contbejel] \
            and lista_SS14[contSS14] != lista_yaws[contyaws] and lista_SS14[contSS14] != lista_bejel[contbejel]:
        contador_NicvsSS14vsYB += 1
        print  "cambio diferente en TPA y no en TEN ni TPE"

    if lista_Nichols[contnichols] != "M" and lista_SS14[contSS14] != "M" and lista_yaws[contyaws] != "M" and lista_bejel[contbejel] != "M" and \
        lista_Nichols[contnichols]!= lista_SS14[contSS14] and lista_Nichols[contnichols] != lista_yaws[contyaws] and lista_Nichols[contnichols] != lista_bejel[contbejel] \
            and lista_SS14[contSS14] != lista_yaws[contyaws] and lista_SS14[contSS14] != lista_bejel[contbejel] and lista_yaws[contyaws] != lista_bejel[contbejel]:
        contador_NicvsSS14vsYvsB += 1
        print  "cambio diferente en todasssss___________________________________-.-.-.-.-.-.-.-.-.-.-.-.-.-.-.-.-.-.-.-.-.-.-"

    return contadorYvsB, contadorNichols_SS14, contador_intersubsp_YBvsSIF, contador_intersubsp_YvsBvsSIF, contador_intersubsp_TPAvsTENTPE,\
           contador_NicvsSS14vsYB, contador_NicvsSS14vsYvsB


def ls(ruta='/home/marta/Documentos/Documentos/Doctorado/MLST_analisis_continuacion/script_SNPs/check_genes_candidates/analisis_script4b_contadores/fastas'):
    ficheros = listdir(ruta)
    return ficheros

def main():
    lista_ficheros= ls()
    for i in lista_ficheros:
        nomfichero = str(i)
        secuencias = abrir_fichero(nomfichero)
        dicc_seq = crear_diccionario(secuencias)
        numseq = len(dicc_seq)
        seq_ref, ref = obtener_ref(dicc_seq)
        dicc_cambios = anotar_cambios()
        dicc_cambios_final, num_cambios_total = procesar_seq(dicc_seq, seq_ref, dicc_cambios)
        dicc_cambios2 = anotar_cambios2()
        dicc_cambios_final2 = procesar_seq_2(dicc_seq, seq_ref, dicc_cambios2)
        diccionario_subespecies_cambi = diccionario_final_bien(dicc_cambios_final, dicc_cambios_final2, numseq)
        contar_cambios_interintra(diccionario_subespecies_cambi, nomfichero)


if __name__ == '__main__':
    main()
```
