## Supplementary_Files for "A new typing scheme demonstrates high discriminatory power for *Treponema pallidum* subspecies": Supplementary_File_3.py

```
#Marta Pla Diaz, Jun 21 (2018), for python2.7


#Execution
#To execute the program, you only need to have the script in the directory where the fastas with the SNP alignments of the genes (or genomes) you want to haplotype are located.
#For each alignment, it will generate two files: one with the SNP alignment and the strains that can be haplotyped, so you can later build the tree with iqtree and visualize the separated haplotypes.
#The other file is a .txt file that contains the reference strain used throughout the program, as well as the list of strains that could not be haplotyped because their SNPs are not informative for this purpose.

from os import listdir

def abrir_fichero(nombre_fichero):
    secuencias = []
    fich = open(nombre_fichero, 'r')
    seq = ""
    for line in fich:
        if line.endswith('\n'): 
            line = line[:len(line) - 1]
        if line.endswith('\r'): 
            line = line[:len(line) - 1]
        if line.endswith(" "):
            line = line.replace(" ", "")
        if line.startswith(">"):
            if seq != "":
                secuencias.append(seq)
                seq = ""
                secuencias.append(line)
            else:
                secuencias.append(line)
        else:
            seq = seq + line
    secuencias.append(seq)
    return secuencias


def crear_diccionario(lista):
    diccionario = {}
    contador = 0
    while contador < len(lista):
        if lista[contador].startswith(">"):
            new_clave = str(lista[contador].replace(">", ""))
            diccionario[new_clave] = ""
            clave = new_clave
            contador = contador + 1
        else:
            diccionario[clave] = lista[contador]
            contador = contador + 1
    return diccionario

def obt_ref(dicc_seq):
    dicc_comp_hech = {}
    dicc_choseref = {}
    contador = 0
    contador_N = 0
    for clave in dicc_seq:
        valor = dicc_seq[clave]
        while contador < len(valor):
            if valor[contador] == "N":
                contador_N += 1
            contador +=1
        dicc_choseref[clave] = contador_N
        contador = 0
        contador_N = 0
    clave_ref = min(dicc_choseref, key=dicc_choseref.get)
    print clave_ref
    valor_ref =dicc_seq[clave_ref]
    dicc_comp_hech[clave_ref] = valor_ref
    return dicc_comp_hech,clave_ref

def comparar_dicc(dicc_seq, dicc_comp_hech):
    lista_news = []
    lista_nohaplotipos = []
    for clave1 in dicc_comp_hech:
        valor1 = dicc_comp_hech[clave1]
        for clave2 in dicc_seq:
            valor2 = dicc_seq[clave2]
            contador = 0
            while contador < len(valor1):
                base1 = valor1[contador]
                base2 =  valor2[contador]
                if base1 != base2 and base1 != "N" and base2 != "N":
                    contador += 1
                    if clave2 not in lista_news:
                        lista_news.append(clave2)
                else:
                    contador += 1
            if valor1 == valor2:
                lista_news.append(clave2)
            if clave2 not in lista_news:
                if clave2 not in lista_nohaplotipos:
                    lista_nohaplotipos.append(clave2)
    for i in lista_news:
        dicc_comp_hech[i] = dicc_seq[i]
    return dicc_comp_hech, lista_nohaplotipos

def crear_alineamiento(dicc_final_haplotipos, nomfichero):
    nomfichero = nomfichero.replace(".fas", "_haplotiposfinales.fas")
    ficheronuevo = open(nomfichero, "w")
    for clave in dicc_final_haplotipos:
        ficheronuevo.write(">")
        ficheronuevo.write(clave)
        ficheronuevo.write("\n")
        ficheronuevo.write(dicc_final_haplotipos[clave])
        ficheronuevo.write("\n")
    ficheronuevo.close()

def crear_sequencias_no_haplotipos(lista_nohaplotipos, nomfichero, clave_ref):
    nomfichero = nomfichero.replace(".fas", "_cepas_NOclasificadas.txt")
    ficheronuevo = open(nomfichero, "w")
    ficheronuevo.write("Cepa utilizada como referencia para comparar: ")
    ficheronuevo.write(clave_ref)
    ficheronuevo.write("\n")
    ficheronuevo.write("Listado cepas no clasificadas: ")
    ficheronuevo.write("\n")
    for i in lista_nohaplotipos:
        ficheronuevo.write(i)
        ficheronuevo.write("\n")
    ficheronuevo.close()

def ls(ruta = '.'):
    return listdir(ruta)

def main():
    lista_archivos = ls()
    for i in lista_archivos:
        if i.endswith(".fas"):
            nomfichero = str(i)
            secuencias = abrir_fichero(nomfichero)
            dicc_seq = crear_diccionario(secuencias)
            dicc_comp_hech, clave_ref = obt_ref(dicc_seq)
            dicc_final_haplotipos, lista_nohaplotipos = comparar_dicc(dicc_seq, dicc_comp_hech)
            crear_alineamiento(dicc_final_haplotipos, nomfichero)
            crear_sequencias_no_haplotipos(lista_nohaplotipos, nomfichero, clave_ref)

if __name__ == '__main__':
    main()
```
